# Genomic analysis reveals shared genes and pathways in human and canine angiosarcoma

**DOI:** 10.1101/570879

**Authors:** Kate Megquier, Jason Turner-Maier, Ross Swofford, Jong-Hyuk Kim, Aaron L. Sarver, Chao Wang, Sharadha Sakthikumar, Jeremy Johnson, Michele Koltookian, Mitzi Lewellen, Milcah C. Scott, Ashley J. Graef, Luke Borst, Noriko Tonomura, Jessica Alfoldi, Corrie Painter, Rachael Thomas, Elinor K. Karlsson, Matthew Breen, Jaime F. Modiano, Ingegerd Elvers, Kerstin Lindblad-Toh

## Abstract

Angiosarcoma is a highly aggressive cancer of blood vessel-forming cells with high fatality and few effective treatment options. It is both rare and heterogenous, making large, well powered genomic studies nearly impossible. In dogs, angiosarcoma is common, with breeds like the golden retriever carrying heritable genetic factors that put them at very high risk. If the clinical similarity of canine and human angiosarcoma reflects shared genomic etiology, dogs could be a critically needed model for advancing angiosarcoma research. We assessed the genomic landscape of canine angiosarcoma via whole exome sequencing (47 golden retriever angiosarcomas) and RNA sequencing (74 angiosarcomas from multiple breeds). The predominant mutational signature was the age-associated deamination of cytosine to thymine, and somatic coding mutations occurred most frequently in the tumor suppressor *TP53* (59.6% of cases) as well as two genes in the PI3K pathway: the oncogene *PIK3CA* (29.8%) and its regulatory subunit *PIK3R1* (8.5%). We compared the canine data to human data recently released by The Angiosarcoma Project, and found the same genes and many of the same pathways significantly enriched for somatic mutations, most notably protein kinases, glycoproteins, fibronectin Type III domains, EGF-like domains, and cell adhesion proteins such as cadherins. As in human angiosarcoma, *CDKN2A/B* was recurrently deleted and *VEGFA, KDR, and KIT* recurrently gained. Canine angiosarcoma closely models human angiosarcoma on a genomic level, and is a powerful tool for investigating the pathogenesis of this devastating disease.

## Introduction

Angiosarcoma is an aggressive cancer of blood-vessel forming cells, associated with poor survival times (1–3). There is an unmet need for new diagnostics and therapies. However, the rarity of this cancer in humans (approximately 0.01% of all cancers) (4,5) has limited large-scale genomic studies so far. Canine angiosarcoma (called hemangiosarcoma in veterinary medicine) is a relevant clinical model for understanding the pathophysiology of human angiosarcomas. The human and canine diseases share many clinical similarities, and the disease is common in dogs, occurring in some breeds (notably the golden retriever) with a frequency up to 20% (6), meaning that using canine angiosarcoma as a model for human disease would yield sample sets of a magnitude inaccessible using human data alone. However, for dogs to be an effective model of this disease in the era of precision medicine, detailed genomic characterization of canine angiosarcoma must be undertaken, and the results directly compared to existing and emerging genomic data from human angiosarcomas.

Angiosarcomas can form anywhere in the vasculature. In human patients, they most commonly occur in the skin of the head, neck, and scalp, the breast, the extremities, and less frequently in the liver, right auricle of the heart, bone, and spleen (7). Prognosis is poor, with metastatic disease occurring in approximately 50% of cases (8), and a median overall survival time of approximately 50 months for local disease and 10 months in metastatic cases (9). Treatment involves surgical resection with wide margins, plus or minus radiation therapy, as well as adjuvant chemotherapy in metastatic disease (3). Many angiosarcomas are initially sensitive to doxorubicin, paclitaxel, or targeted agents, but resistance to these therapies is virtually inevitable (3).

While most cases of angiosarcoma occur without known cause, there are several known risk factors. These tumors can arise secondary to radiation therapy for other cancers or chronic lymphedema (3). Other known risk factors include UV irradiation, given the typical locations of cutaneous angiosarcomas on the head and neck (10–12), as well as occupational exposure to vinyl chloride (13), arsenicals, and use of anabolic steroids (14). Genetically, angiosarcoma is associated with familial syndromes including Li-Fraumeni syndrome (*TP53* mutations) (15) and Klippel-Trenaunay syndrome (*PIK3CA* mutations) (16). However, these syndromes do not solely cause angiosarcoma (7), and are not present in the majority of cases.

Canine angiosarcoma is the histopathological equivalent of the human disease (17), and follows a similar, aggressive clinical course. In dogs, the most common locations are the spleen, right auricle of the heart, liver, and skin or subcutaneous tissue (18,19). The difference in distribution of locations from the human disease is likely due in part to the lack of secondary cases in dogs. Treatment protocols similarly involve wide surgical resection, followed by adjuvant chemotherapy. Survival times are short - a median of 4-6 months after surgical resection with adjuvant chemotherapy, with a 1 year survival rate of 10% (20–22). Biological risk factors have not yet been identified in dogs. Genetically, dog breeds display differential predisposition to specific cancers, indicating that there are heritable risk factors that have become common as a result of inbreeding based on selection or drift. In a previous genome-wide association study, we identified several loci significantly associated with the risk of disease in the golden retriever (23).

Recent targeted next-generation sequencing of human angiosarcomas has begun to reveal the somatic mutational spectrum of this disease. It has so far proven to be fairly heterogeneous - no pathognomonic mutations or copy number aberrations occur in all cases, and tumors in different primary locations or with different underlying etiologies have genomic differences. *TP53* and genes in the MAPK pathway are frequently mutated (24), and recurrent mutations in *PLCG1* and *PTPRB* are common, particularly in secondary angiosarcomas (24,25). Although the PI3K pathway is activated in some human angiosarcomas (26,27), *PIK3CA* mutations have so far not been commonly reported in human angiosarcoma studies (25,28). Genes in the VEGF pathway are frequently gained or amplified, including *VEGFA*, and *KDR* (24,27). The tumor suppressor *CDKN2A* is frequently deleted, while the *MYC* oncogene is frequently amplified, most commonly in radiation-associated tumors (24,27).

In dogs, a recent whole-exome sequencing study of a small cohort of 20 angiosarcomas showed that the top recurrently mutated genes were *PIK3CA* and *TP53* (29). Earlier candidate gene studies of canine angiosarcomas have reported mutations in *TP53 (30)*, *PTEN (31)*, and *PDGFRA* and *PDGFRB (32)*. An analysis of somatic copy number aberrations in visceral angiosarcomas from five breeds found that *VEGFA* showed frequent copy number gain, while *CDKN2A* was frequently deleted (33). *MYC* copy number gain was infrequent, likely reflective of the fact that secondary angiosarcomas are not seen in canine patients (33). While these earlier studies provide clues as to the genetic features of canine angiosarcoma, there is very little of the genome-wide data needed for comprehensive comparison to human disease.

In order to assess the potential utility of canine angiosarcoma as a model for human angiosarcoma at the molecular level, we performed the largest exome sequencing study of canine angiosarcoma to date, and complemented our exome data with oligonucleotide array comparative genomic hybridization (oaCGH) copy number data and RNA sequencing (RNA-seq) data in partially overlapping cohorts of canine angiosarcoma cases. We then performed comparative analyses of our results with those released by The Angiosarcoma Project (AP) direct-to-patient initiative. In this way, we have created a detailed genomic profile of this cancer in the golden retriever breed, and begun vetting this canine cancer as a comparative model for human angiosarcoma, and potentially other tumors.

## Materials and Methods

Additional Materials and Methods can be found in the **Supplementary Materials and Methods**, and a workflow in **Figure S1**.

### Canine angiosarcoma sample collection for exome sequencing

Samples were obtained as part of necessary diagnostic procedures with owner consent. DNA from tumor tissue and whole blood was collected from 47 golden retrievers with visceral angiosarcoma of the spleen, heart, or liver (See **SI** for sample locations).

### Sample preparation

DNA was extracted and sequencing libraries prepared using the Kapa Hyper Prep Kit (**Table S2**). For 66 samples, additional cycles of PCR were required to obtain sufficient DNA for exome capture, while libraries from 28 samples were re-constructed from source DNA using the standard protocol (See **SI**).

### Exome capture

The Roche Nimblegen SeqCap-EZ capture canine exome (120705_CF3_Uppsala_Broad_EZ_HX1), which targets approximately 49 Mb of exonic and regulatory sequence, was used for hybrid exome capture, following the manufacturer’s protocol.

### Sequencing and read alignments

The barcoded exome-captured libraries were multiplexed in pools of 8, and sequenced on the Illumina HiSeq 2500 to a target depth of 60x in the tumor and 30x in the normal, reaching a mean depth of 78x in the tumor and 63x in the normal. Reads were aligned to the CanFam3.1 reference genome using BWA. PCR duplicate reads were flagged using the Picard tool MarkDuplicates. Following the Genome Analysis Toolkit (GATK) Best Practices, we then performed Base Quality Score Recalibration (BQSR) using a set of approximately 19 million known canine germline variant locations.

### Somatic variant calling

Somatic mutations were then called using MuTect2, using the default settings, with the addition of the *--dontUseSoftClippedBases* option, which was added to avoid calling a large number of artifactual indels in our FFPE-preserved samples (See **SI**). In order to further refine our set of somatic variant calls, and to avoid artifacts, we also called variants using the GATK4 version of Mutect2, and kept only the consensus of calls which passed in both the GATK3 and GATK4 versions. Both sets of calls were filtered using a “panel of normals” created using all 47 normal (germline) samples. Calls were further filtered to exclude oxidation artifacts, and were removed if they overlapped locations with known germline variants. Variants were further filtered if the position had low coverage (defined as a read depth less than 20 in the normal, less than 40 in the tumor, or less than 4 alternate allele reads in the tumor), or had excessive read depth (greater or equal to mean read depth + 5 x standard deviation), to filter out potential alignment errors.

### Variant annotation

Variants were annotated using the variant effect prediction program SnpEff v4.2. Where multiple effects were predicted for a single variant, the most damaging predicted effect was selected. Coding variants were analyzed using the *smg* function in Genome MuSiC 0.4 to determine which genes were significantly mutated above the background rate. The *merge-concurrent-muts* option was applied to count multiple mutations in the same gene within a sample as a single mutation, and a false discovery rate (FDR) threshold of 0.1 using the convolution test (CT) was applied.

### Canine mutational signature discovery

Mutational signatures, considering point mutations and their genomic context, were extracted using a Bayesian non-negative matrix factorization (NMF) algorithm. The discovered signatures were compared to known signatures in COSMIC (34), as well as those reported in the literature for human angiosarcomas (35). The overall landscape of mutations was plotted for different groupings of samples using the SomaticSignatures package.

### RNA-sequencing of canine angiosarcomas

Seventy-six snap-frozen samples were obtained from seventy-four dogs with angiosarcoma (42 golden retrievers and 32 dogs from 12 other breeds or from mixed breeding) and ten dogs with splenic hematomas (Kim, *et al.*, Cancer Res, 2018, submitted manuscript). RNA-seq data from forty-seven of the angiosarcoma tissues had been published in our previous studies (23,36). RNA-seq libraries were generated as described previously (36).

### Somatic copy number detection in canine angiosarcomas

Analysis of somatic copy number aberrations (SCNAs) in our canine whole exome sequencing data was limited by the inclusion of both frozen and formalin-fixed (FFPE) samples. The FFPE samples appeared to have an increased number of amplifications and deletions when compared with the frozen samples, possibly as a result of DNA fragmentation during the fixation process (37). While we were able to partially account for this using GC correction (38), we were not able to completely remove the bias between the two groups (**Figure S6**). Hence, we instead used oaCGH data to interrogate copy number changes in a cohort of 69 golden retrievers with angiosarcoma tumors, of which 28 were also included in the exome sequencing cohort. SCNAs were called from oligonucleotide array comparative genomic hybridization (oaCGH), as described previously (33), using a ~180,000-feature microarray (Agilent Technologies) with approximately 26 kb resolution throughout the dog genome.

### Accessing human data from The Angiosarcoma Project for comparative analysis

The results included here are based on the use of data from The Angiosarcoma Project (AP, https://ascproject.org/), a project of Count Me In (https://joincountmein.org/), downloaded October, 2018. We compared the somatic mutations and SCNAs present in our cohort of canine angiosarcomas to the results reported by the AP, a direct-to-patient sequencing project run by the Count Me In initiative (JoinCountMeIn.org). Mutations and SCNAs derived from whole exome sequencing data from 48 samples from 36 patients were downloaded from CBioPortal, through the “Mutated Genes” or “CNA Genes” charts on the project main page. CBioPortal applied the following filters: somatic mutations were non-silent coding mutations which occurred in two or more samples, or that occurred in one sample but in a known cancer gene. Copy number aberrations were reported if they occurred in at least one sample in a known cancer gene. Primary tumor locations included 19 breast tumors; ten head, face, neck, or scalp (HFNS) tumors; two pulmonary tumors; and one tumor each in bone, heart, spleen, bladder, and abdomen.

### Pathway analysis in canine and human data

Pathway analysis was performed on the canine and human mutational data using the DAVID Functional Annotation Tool. For the overall enrichment analyses, all genes with somatic mutations were entered. Functional annotation clustering was performed using the default options, using the Benjamini-Hochberg method to control the false discovery rate. Analysis was performed independently for the canine data and the human Angiosarcoma Project data, mapping genes from both species to *Homo sapiens* to avoid confounding due to differences in gene annotation between the species. Enriched clusters were ranked so that the top three pathways for each of the top 10 clusters were compared between dogs and humans.

## Results

### Somatic SNVs and indels

#### TP53 and PIK3CA are commonly mutated in canine angiosarcomas

The significantly mutated genes (SMGs) in the canine angiosarcomas contained well-known cancer genes, including *TP53*, as well as two genes in the PI3K pathway. Overall, seven genes were significantly mutated (Table 1, Figure 1). Tumor suppressor *TP53* was most frequently mutated (28/47 cases, 59.5%), with all 28 cases carrying at least one mutation affecting the DNA binding domain (Figure 2). Oncogene *PIK3CA* (14/47, 29.8%) and its regulatory subunit *PIK3R1 (*4/47, 8.5%) were both mutated. Ten of the fourteen cases with *PIK3CA* mutations had a mutation at amino acid position 1047, a hotspot frequently mutated in many types of human cancers (39) (Figure 2). The remaining four SMGs were *ORC1* (4/47, 8.5%), *RASA1* (4/47, 8.5%), *ARPC1A* (3/47, 6.3%), and *ATP5PD* (2/47, 4.2%). Overall, 33 genes that were mutated at least once in our dataset are annotated as likely causal in the COSMIC Cancer Gene Census, including *TP53*, *PIK3CA*, and *PIK3R1* (**Table S3**).

**Table 1.**
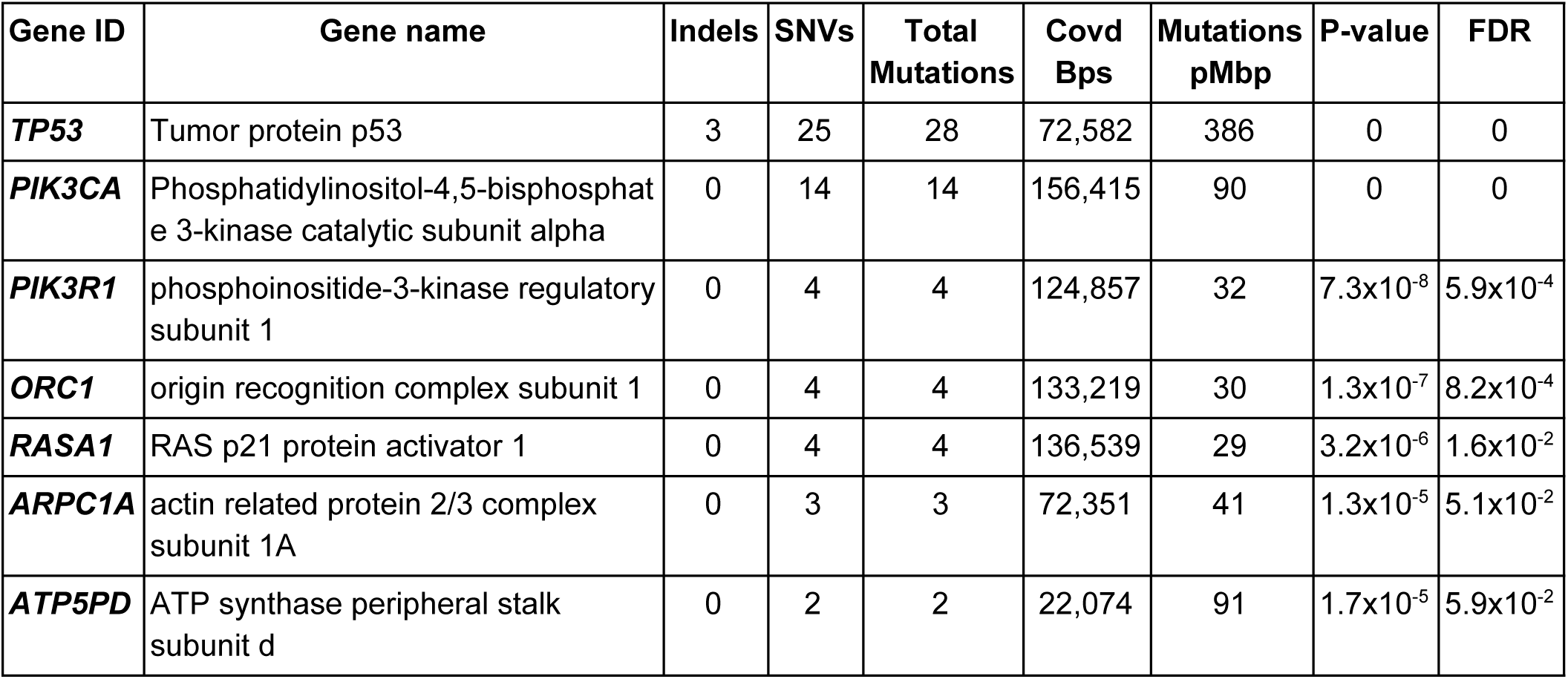
Significantly mutated genes in golden retriever angiosarcoma tumors. Significantly mutated genes with an FDR < 0.1, calculated using Genome MuSiC. SNVs: single nucleotide variants, Covd Bps: number of basepairs with adequate coverage in gene, Muts pMbp: mutations per megabase.

**Figure 1.**
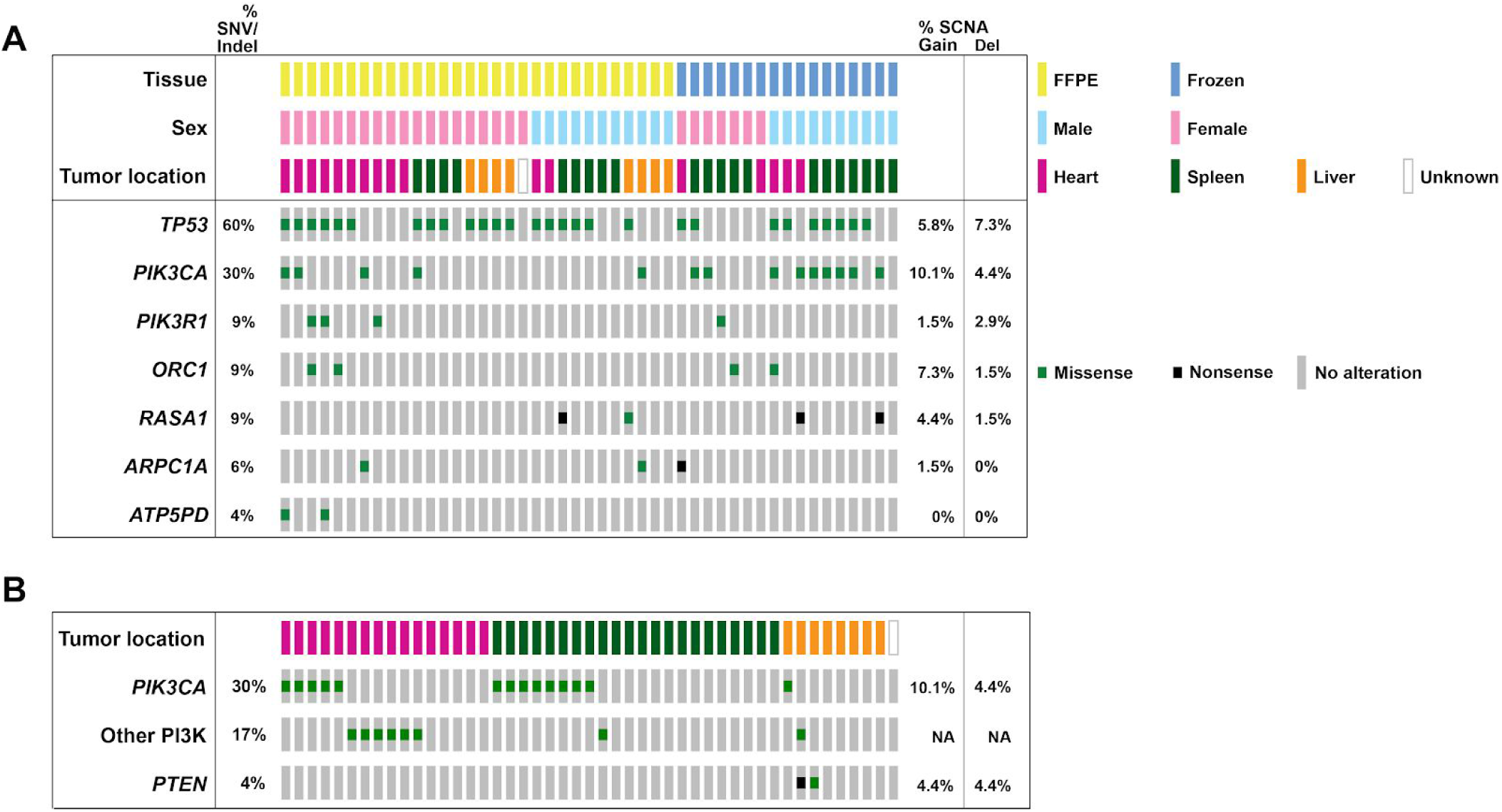
Per-sample annotation of metadata. Including: A) somatic mutations in the seven significantly mutated genes B) somatic mutations in the PI3K pathway and tumor location. ‘Other PI3K’ includes mutations in *PIK3CB*, *PIK3C2G*, *PIK3C3*, *PIK3R1*, and *PIK3R5*). SNV, single nucleotide variant; Indel, insertion/deletion; SCNA, somatic copy number aberrations. SCNA frequencies are calculated based on oaCGH data from 69 canine angiosarcoma samples.

**Figure 2.**
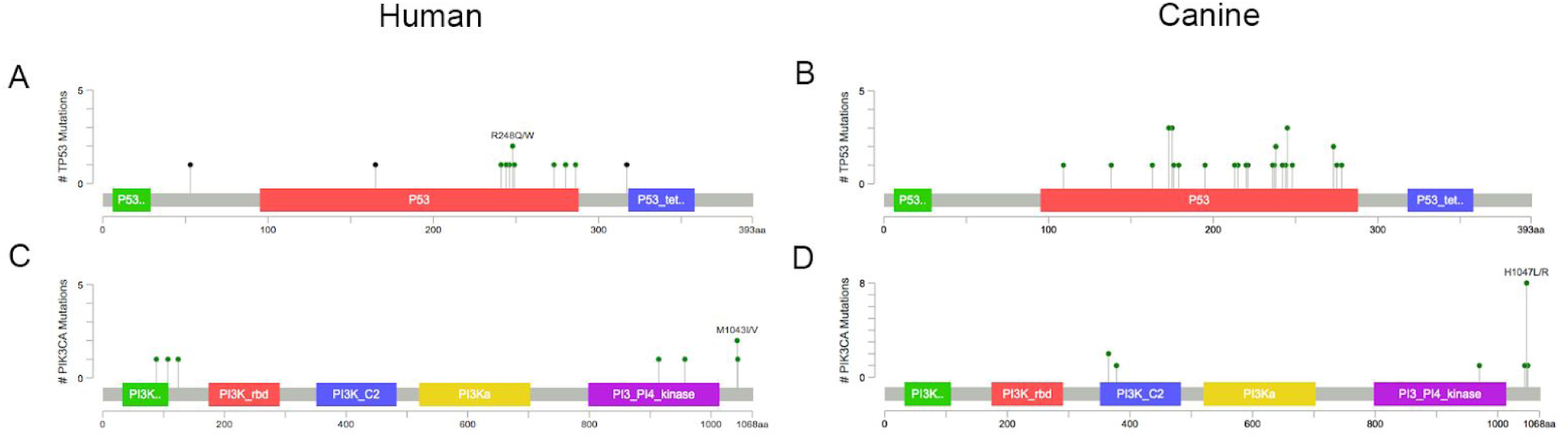
Lollipop plots showing the positions of the TP53 (A and B) and PIK3CA (C and D) mutations in the human (left) and canine (right) data. Canine mutational locations were lifted over to the human hg19 reference genome using the LiftOver tool.

#### Comparison with top mutated genes in human angiosarcoma

The most commonly mutated genes in the human AP data were *TP53* (11/36 patients, 31%), *KDR*, *RYR2*, *TTN*, *LRP2*, *PKHD1*, and *ABCA13* (8/36 patients affected each, 22%), followed by *PIK3CA, RYR1, ASXL3, MYH14, FLG, MUC16* (in 7/36 patients each, 19%). Two AP cases (6%) also had *RASA1* mutations. In the canine cohort, many of these same genes were mutated, but at much lower frequencies: *KDR, RYR2, LRP2, ABCA13, RYR1, ASXL3*, and *MUC16* each had nonsynonymous mutations in one case (2%), *TTN* had nonsynonymous mutations in five cases. No nonsynonymous mutations were found in *PKHD1*, *MYH14*, or *FLG*.

#### Canine RNA-seq analysis validates exome mutations and extends findings to additional breeds

We validated our somatic mutation calls from exome sequencing using RNA-seq data from a partially overlapping set of dogs. Fifteen golden retrievers from the exome cohort with mutations in SMGs were included in the RNA-seq cohort, with a total of 28 SMG mutations among them. Of these sites, we validated 15/28 (53.5%) mutations, with low variant allele frequency (VAF), nonsense-mediated decay, and tumor heterogeneity likely contributing to the 46.5% that were not replicated (**Table S4**).

The RNA-seq cohort (n = 76 tumors obtained from 74 dogs from 12 breeds), also confirmed that the *TP53* and PI3K mutations were not breed-specific, but were present across many different breeds. In addition to the 15 golden retrievers also included in the exome cohort, somatic mutations corresponding to those observed in the exome cohort were noted in *TP53* (n = 16, 27.1%, 5 breeds), *PIK3CA* (n = 12, 20.3%, 7 breeds, including mixes), and *PIK3R1* (n = 4, 6.7%, 4 breeds, **Table S5**).

#### Comparison of mutational burden by tumor location in canine and human data

The canine tumors had fewer nonsynonymous mutations than the human tumors, while the mutational burden varied significantly by tumor location in human cases. In the canine cohort, there were a median of 22 nonsynonymous coding mutations per sample (range 1-149, **Table S1**). There was no difference in the total number of nonsynonymous mutations between heart, splenic, and liver tumors (p_ANOVA_ = 0.04, Tukey HSD test not significant). In the AP human data, there was a significant difference in the median number of mutations by tumor location, with head, face, neck, and scalp (HFNS) tumors having a higher mutational burden than breast or visceral tumors (p_anova_ < 0.0001)(40). The distribution of median mutational burden in the human tumors was: HFNS (n = 12, median = 398.5), breast (n = 26, median = 32, outlier with 344 mutations removed), visceral (n = 7, median = 41).

#### Differential patterns of mutations found in some tumor locations

The pattern of nonsynonymous mutations varied more when we analyzed only the PI3K gene family, with the number of mutations in this gene family varying by tumor location in both dogs and humans. In the canine cases, a slightly higher proportion of heart tumors (11/16, 68.8%) compared to splenic tumors (9/22, 40.9%) had PI3K alterations, although this difference was not significant (p_chi-sq_=0.09). Liver tumors had slightly fewer overall alterations in the PI3K gene family (3/8, 37.5%), and were significantly less likely to have *PIK3CA* mutations (1/8, 12.5%) than heart tumors (p_chi-sq_=0.009). In the human AP data, breast tumors were the only location to carry *PIK3CA* mutations (n=10 samples, 7 patients).

#### Comparative pathway analysis reveals similarities between angiosarcoma in dogs and humans

Pathway analysis of all genes with non-silent coding somatic mutations in the canine dataset, and the CBioPortal-filtered human dataset (n_canine_=951, n human =1934) revealed striking similarities between the two species, despite the difference in the distribution of tumor locations, and the absence of secondary tumors in the canine cohort (**Table 2, Table S6, Table S7**). Comparison of the top ten most enriched pathways in both species revealed that both were enriched for glycoproteins (p_human_ = 1.0 × 10^−52^, p_canine_ = 5.6 × 10^−15^), fibronectin type III domains (p_human_ = 9.2 × 10^−23^, p_canine_ = 2.5 × 10^−13^), cadherins and adhesion proteins (p_human_ = 3.5 × 10^−16^, p_canine_ = 1.1 × 10^−3^), epidermal growth factor domains (p_human_ = 1.0 × 10^−20^, p_canine_ = 2.1 × 10^−5^), ATP binding domains (p_human_ = 5.4 × 10^−18^, p_canine_ = 6.2 × 10^−8^), and protein kinases (p_human_ = 7.4 × 10^−17^, p_canine_ = 5.8 × 10^−3^). Pathway analysis of the overlapping genes mutated in both cohorts (n = 267, **Table S8**), as well as the union of all mutated genes in both cohorts (n = 2617) yielded similar results (**Table S9**).

**Table 2.**
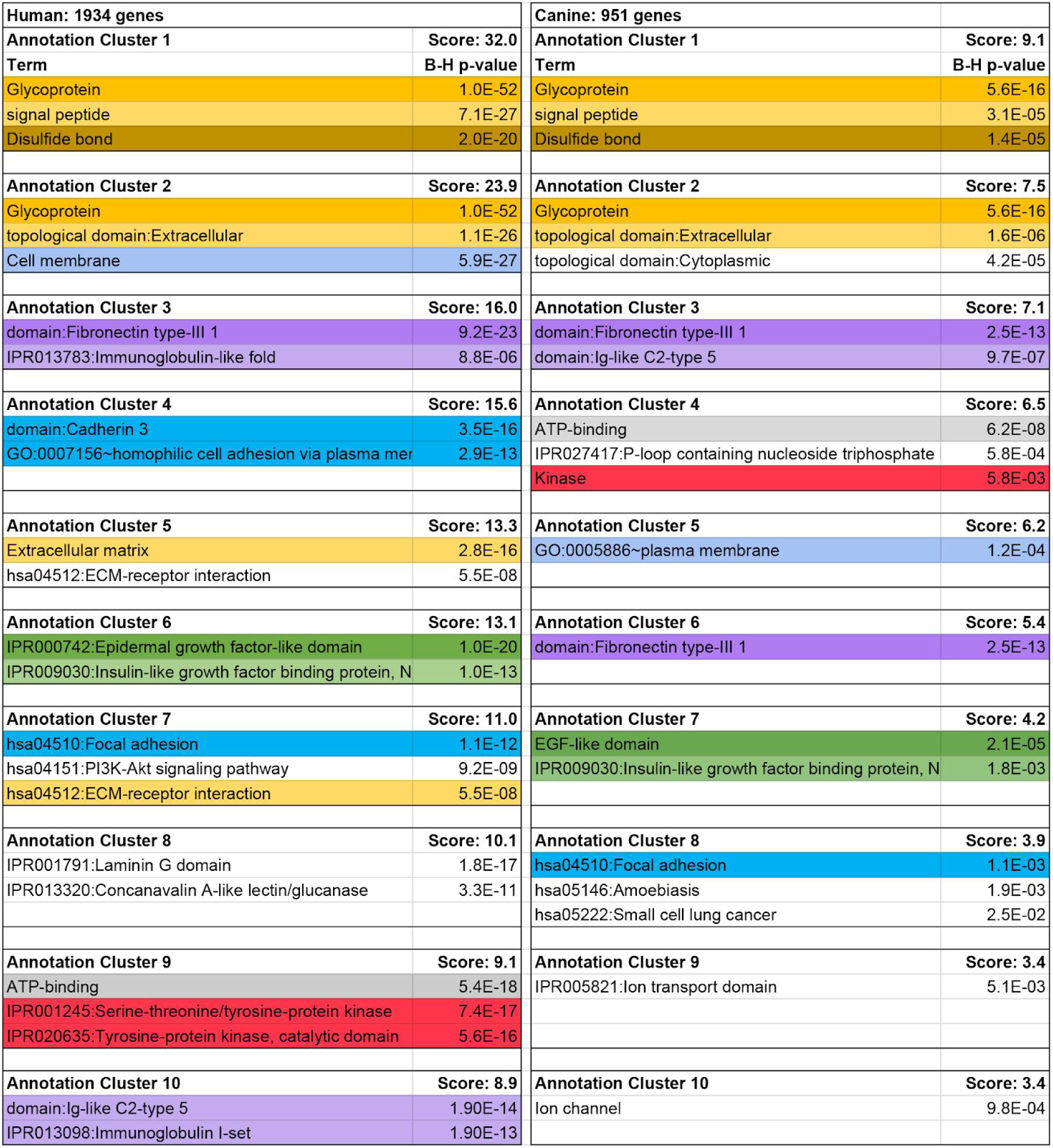
Comparison of the top 10 enriched functional annotation clusters in the human and canine data. Functional annotation clustering was performed in DAVID, and the top three pathways in each of the top 10 enriched clusters were compared between the canine and human data. Pathways that are enriched in both species are highlighted in matching colors. Human data contained non-silent coding mutations which occurred in two or more samples, or that occurred in one sample but in a known cancer gene. Canine data included all non-silent coding mutations.

#### Shared pathways and gene families previously reported in human angiosarcoma literature

Several of the less significantly enriched pathways and gene families in both dogs and humans have been previously reported to be affected in the human angiosarcoma literature. Phospholipase C (PLC) genes were enriched in the 267 genes mutated in both species (p = 4.1 × 10^−3^). One case in our canine cohort had a mutation in *PLCG1*, a gene recurrently mutated in human angiosarcomas (25), while eight cases (17%) had mutations in other PLC genes. In the human AP cohort, 14 (29.1%) samples had mutations in 7 PLC genes, including *PLCG1* (n=8, 16.7%, Table 3). Both the canine and human datasets were separately enriched for protein tyrosine phosphatases (p_human_= 7.3 × 10^−5^, p_canine_= 4.8 × 10^−3^). While none of the canine cases had somatic mutations in the *PTPRB* gene, a gene recurrently mutated in human angiosarcomas (25), seven cases (14.9%) harbored a mutation in one of seven protein tyrosine phosphatase genes. Sixteen protein tyrosine phosphatases were mutated in sixteen samples in the human AP tumors (Table 3). Finally, both the canine and human datasets were enriched in LDL receptors (p_human_ = 0.017,, p_canine_= 0.022, Table 3). While mutations in this family of receptors have not been widely reported in human angiosarcomas, they have recently been shown to play a role in a variety of cancers (41), and *LRP2* is one of the most frequently mutated genes in the AP dataset.

**Table 3.**
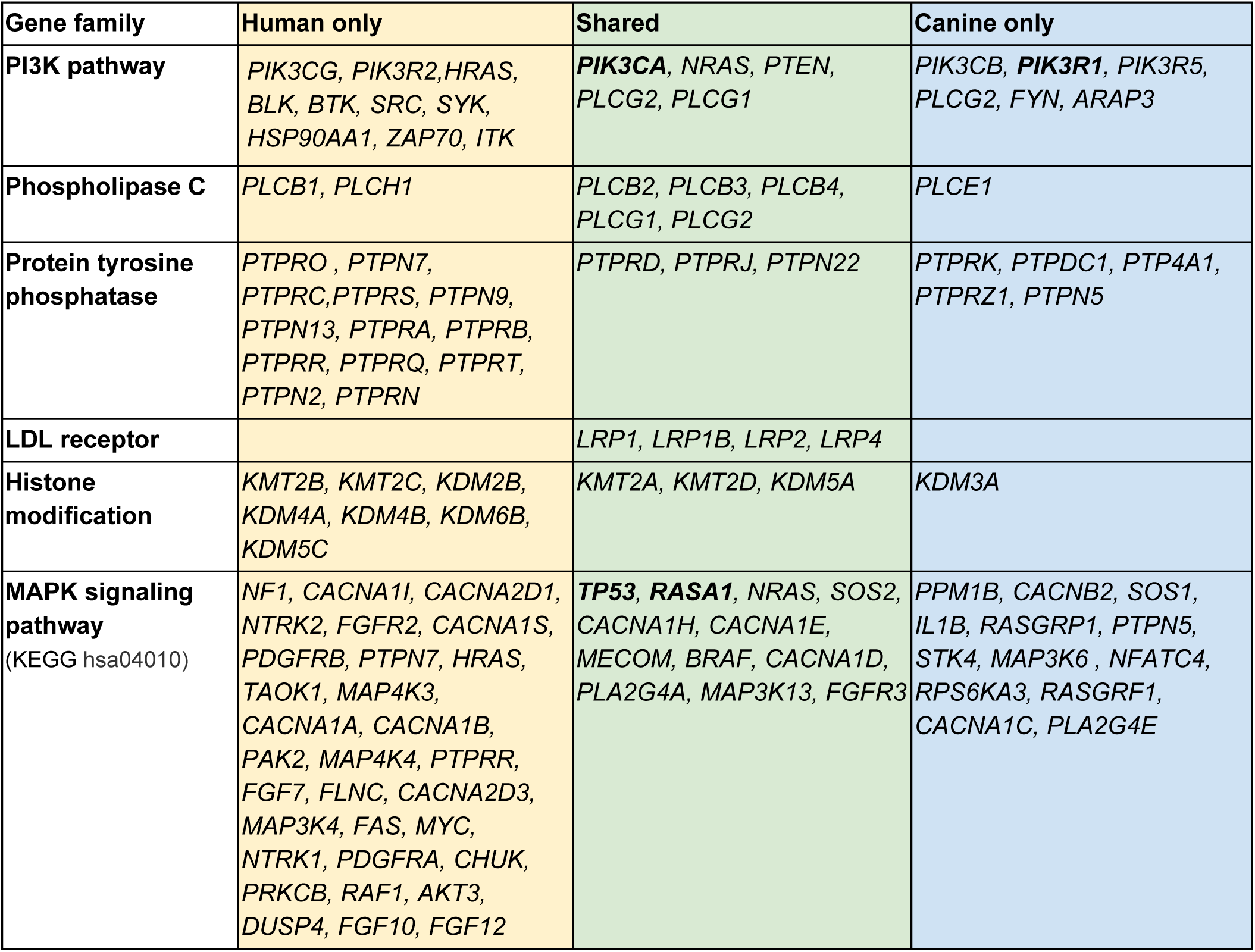
Overlap of mutated genes between canine and human angiosarcoma tumors in several gene families and pathways. Significantly mutated genes bolded. Note: Human data contained non-silent coding mutations which occurred in two or more samples, or that occurred in one sample but in a known cancer gene. Canine data included all non-silent coding mutations.

We noted mutations in 25 genes in the MAPK pathway in the canine cohort, including the SMGs *TP53* and *RASA1*. In the human AP cohort, 43 genes in the MAPK pathway were mutated. Twelve genes in the MAPK pathway were mutated in both the human and the canine cohorts (Table 3). However, neither the human nor the canine datasets were enriched in the MAPK pathway in the DAVID analysis. Similarly, although we found mutations in histone modifying genes, consistent with earlier human studies (42), we saw no significant enrichment on a pathway level in either the canine and human cohorts (Table 3).

#### Mutational signature of aging is present across all canine tumors

Analysis of mutational signatures in the canine tumors, which capture the mutational landscape of tumors and are shaped by both genetic factors and environmental exposure revealed similarities with human angiosarcoma. We analyzed the signatures of trinucleotide mutational frequencies present in golden retriever angiosarcomas, and found a strong signature of mutations arising through spontaneous deamination of methylated cytosines in CpG islands, corresponding to COSMIC signature 1 and generally associated with aging (34) (Figure 3, **Figure S3-S5)**. In addition, we see a very faint signature not described in COSMIC (Figure 3). The overall mutational landscape was consistent across all canine angiosarcoma tumors, regardless of tumor location or tissue preservation method (**Figures S2 - S4**).

**Figure 3.**
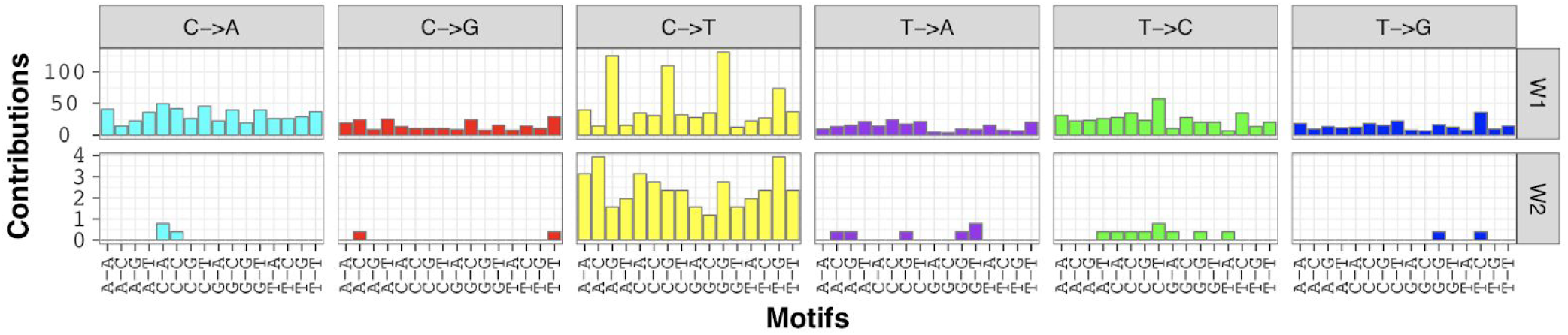
Mutational signatures called using Bayesian NMF in the entire canine angiosarcoma cohort, showing the count of mutations (y-axis) in each trinucleotide context (x-axis). This analysis reveals the COSMIC 1 aging signature (W1), and a faint secondary signature not matching any COSMIC signature (W2).

### Somatic copy number aberrations

#### SCNAs in canine angiosarcoma recurrently affect known cancer genes

The genes most recurrently affected by DNA copy number aberrations in the oaCGH data were *VEGFA* showing copy number gain in 19% of cases, *KDR* gain in 22%, *KIT* gain in 17%, and the tumor suppressor *CDKN2A/B*, deleted in 22% (**Table S10**). The *MYC* oncogene showed copy number gain in 9% of cases. Copy number aberrations in the top significantly mutated genes were relatively rare (Figure 1A, **Table S10**).

#### Copy number gains in KDR and KIT are common in both dogs and humans

Comparison of SCNAs in the filtered human AP data and canine oaCGH data revealed recurrent copy number gains in known cancer genes *KDR* and *KIT* in both species (**Table S11**). Copy number gains in *KDR* occurred in approximately 31% of human samples, and 22% of canine samples. KIT was gained in 17% of both human and canine samples.

#### SCNA profiles differ among cases with and without PIK3CA mutations

We examined the SCNA profiles of cases with and without *TP53* and *PIK3CA* mutations in the 28 cases with both exome sequencing and oaCGH data. There were no significant differences in the relative frequency of any given CNA between the CNA profiles of cases with and without *TP53* mutations. However, significant differences were detected between cases with *PIK3CA* mutations and those without (Figure 4), using a two-tailed Fisher’s Exact test and a minimum differential threshold of 25% between the two groups. A region on chromosome 11 at approximately 22.6 Mb, near the putative *UBE2B* (ubiquitin-conjugating enzyme E2B) gene, was deleted in 4/11 cases with *PIK3CA* mutations, and 0/17 without (p < 0.016). Similarly, the *CDKN2B* gene, located distally on chromosome 11 at 41.2Mb, was deleted in 4/11 cases with *PIK3CA* mutations, and 0/17 cases without (p < 0.016). A region on chromosome 24 at 21.2 Mb was gained in 7/17 cases without *PIK3CA* mutation and 0/11 with *PIK3CA* mutation (p < 0.023). This region overlaps the anti-apoptotic *BCL2L1* gene. Broad copy number gains along the length of chromosome 31 were more frequent in cases without *PIK3CA* mutation compared to those with *PIK3CA* mutation.

**Figure 4.**
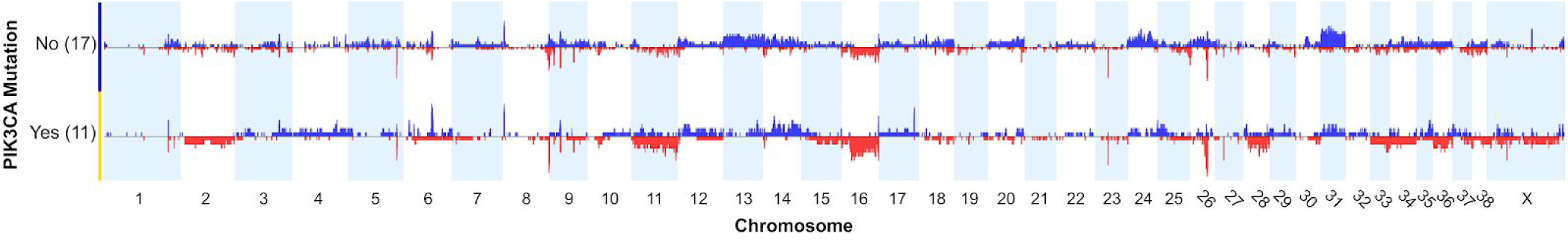
Comparison of DNA copy number aberration profiles between cases with *PIK3CA* mutation and those without. (Thomas, *et al.*, manuscript in preparation)(33)

## Discussion

We have shown through detailed molecular profiling of canine angiosarcoma that the genetic landscape of these tumors is similar between dogs and humans, complementing their similarity in clinical presentation and outcome. Our findings have important implications for comparative oncology, as the study of canine angiosarcoma as a model for the human disease has the potential to improve our understanding of the pathophysiology of the disease, and to improve treatment and outcomes in both species.

Tumor suppressor *TP53* was the top significantly mutated gene in our canine cohort. The majority of mutations occurred in the DNA binding domain, likely causing loss of function (Figure 2). *TP53* is also the only significantly mutated gene in the human AP data, and it has been frequently reported as mutated in targeted sequencing studies of human angiosarcoma (24).

We found that the PI3K pathway was commonly mutated in both canine and human angiosarcoma. A total of 23 canine tumors (48.9%) had at least one somatic mutation affecting this gene family (Figure 1A). Tumors with a mutation in the PI3K family tended to have only one mutation in this family. The PI3K pathway is one of the most commonly altered pathways in cancer, playing an important role in signal transduction leading to cell proliferation, survival, differentiation, and regulation of metabolism and immunity (43,44). *PIK3CA* is an oncogene (45) that has been shown to be mutated in human glioblastoma, breast, gastric, colorectal, lung, and endometrial cancers (46). Ten of the 14 *PIK3CA* mutations in our canine cohort occurred at amino acid position 1047, a mutational hotspot in many human cancers (39). Mutations within this domain have been shown to increase catalytic activity (46–48).

There were also some potentially important differences in somatic mutations between the two species. In the human data, mutations in *TP53* and *PIK3CA* tended to be mutually exclusive, while we do not see this pattern in the canine tumors. In addition, *PIK3CA* mutations were exclusively found in breast tumors in the human AP data, while we found them to be common in cardiac and splenic tumors in the dogs. Within the canine visceral tumors, we found significantly fewer mutations in *PIK3CA* in liver tumors than in cardiac tumors. These differences in distribution of *PIK3CA* mutations by tumor location may be due to genetic heterogeneity of the cancer, with tumors in different locations activating the PI3K pathway at different points, or relying on alterations in different pathways to affect an essential protein downstream. It is also possible that this difference is artificial, due to the small number of human visceral angiosarcomas currently sequenced, and the small number of liver tumors in the canine cohort.

Another potentially important difference between the two species is that, while copy number gains in *KDR* are common in both species, somatic mutations in this gene were seen in over 20% of human tumors, but only one canine tumor. As the *KDR* receptor is upstream of the PI3K pathway, it is possible that mutations in either may lead to a similar phenotype. We also saw more *VEGFA* gains in our canine cohort (19% vs. 0 in the AP human data), and as *VEGFA* is upstream of *KDR*, this copy number gain may serve a similar role to *KDR* mutations in the canine tumors.

Comparative pathway analysis revealed that both the canine and the human tumors are enriched for somatic mutations in a number of the same protein types, domains, and functional pathways that are important for cancer pathophysiology. Overall the pattern of enrichment points to dysregulated signaling, which occurs in many cancers. While the broad enrichment categories are not specific for this particular type of cancer, the shared distribution of these enrichments in canine and human angiosarcomas points to their underlying similarity.

Several of the pathways and domains enriched for mutations in both the canine and human tumors function in the extracellular matrix, cell-cell interaction, and cellular adhesion. These include glycoproteins, fibronectin type III domains, adhesion proteins, and EGF-like domains. Fibronectin type III domains are protein domains that are important for the interaction of cells with the surrounding extracellular matrix, playing an important role in cell adhesion, migration, and signaling (49). Proteins involved in cellular adhesion were also enriched, including cadherins, loss of which promote tumor invasiveness and metastasis, as well as stimulating cellular proliferation (reviewed in (50)).

Tumors in both species were also enriched for mutations in tyrosine kinases, which are important regulators of cellular growth and division signals and are commonly mutated in cancers. Downstream from these mutations, tumors in both species were enriched for mutations in the protein tyrosine phosphatase gene family. This may suggest an alternate mechanism of tyrosine kinase overactivation, as protein tyrosine phosphatases deactivate tyrosine kinase signaling by dephosphorylating proteins in opposition to kinase phosphorylation (51). Further downstream, phospholipase C proteins play a crucial role in cellular signalling pathways by hydrolyzing phosphatidylinositol 4,5-bisphosphate (PIP2) into the second messengers DAG and IP3, passing on signals from receptor tyrosine kinases (52). Both the human and canine angiosarcomas were enriched for phospholipase C genes, including *PLCG1*, although *PLCG1* mutations were not recurrent in the canine tumors.

The shared enriched pathways between canine and human angiosarcomas provides insight into disease pathogenesis for both species. Tyrosine kinase inhibitors have been effective against angiosarcoma in the clinic, but tumor heterogeneity and the development of resistance have limited their long-term utility. Future investigation of the interaction between these pathways will help to determine the potential for combination therapy targeting multiple of these pathways or a common downstream effector to combat resistance.

A recent study in radiation-induced breast angiosarcomas detected the irradiation signature, as well as the aging signature, and a unique C>T signature (35). Canine angiosarcomas looked very similar, in that we primarily saw the aging signature and low levels of a not-yet reported C>T signature. This weak novel signature bore some resemblance to the signature reported in the human angiosarcomas, including higher levels of C>T mutations at C nucleotides flanked by A-A or A-T, however, there were also differences, such as a high number of mutations flanked by T-G in the dogs, and a low number of these in the human cohort. A larger study will be needed to decipher whether this is a novel angiosarcoma-related signature or whether it represents noise, particularly as it is so faint in the canine data. The lack of the irradiation signature was anticipated as our data did not include any tumors secondary to radiation therapy.

The genes most commonly affected by copy number aberrations are those that have been reported previously in both canine and human angiosarcoma cases - copy number gain of *VEGFA* and *KDR*, and *CDKN2A/B loss*. The *MYC* oncogene, which has been reported amplified in human and canine angiosarcomas, is only rarely gained in our data set and shows no evidence of high-level amplification. This makes sense given that it is more common in radiation-induced tumors, which are not present in this canine cohort.

Our data suggest that visceral canine angiosarcoma can be developed as an excellent model for primary human angiosarcomas of the viscera and potentially the breast. Detailed molecular characterization of canine angiosarcoma revealed many similarities between the canine and human angiosarcomas. This was particularly apparent at the pathway level. Future work should entail analysis of larger sample sizes in order to decipher potential molecular subtypes and to facilitate a more complete comparison between tumors in different locations. An integrated understanding of the interaction between mutations in the many enriched signaling pathways may be useful for determining treatment strategy, for example, the feasibility of combinations of targeted inhibitors or the prevention of convergent resistance. Our data suggest that clinical trials evaluating therapeutic approaches in dogs could also inform human medicine.

## Supporting information

Supplementary methods, discussion, tables, and figures

## Acknowledgments

The authors would like to thank all of the dogs and owners who participated in our research, as well as the veterinarians who collected samples. In addition we would like to thank Count Me In. We would also like to thank Dr. Scott Moroff, Vice-President and Chief Scientific Officer of Antech Diagnostics, for contributing tumor and paired blood samples to our research. This work was funded by American Kennel Club (AKC) Canine Health Foundation (CHF) grants #422 (JFM), 1131 (JFM), and 1889-G (JFM, MB, KLT, EKK), NIH grants R03CA191713 (JFM), P30 CA077598 (NIH Comprehensive Cancer Center Support Grant to the Masonic Cancer Center, University of Minnesota) and R01CA218570 (KM, EKK), NCCF grants DM06CO-003 (JFM) and JHK15MN-004 (JHK), and a Morris Animal Foundation grant D10CA-501 (JFM, MB, KLT). KLT is the recipient of a Distinguished Professor award from the Swedish Research Council. IE is supported by a postdoctoral fellowship from the Swedish Medical Research Council, SSMF. JFM is supported by the Alvin and June Perlman Chair in Animal Oncology at the University of Minnesota. MB is supported in part by the Oscar J. Fletcher Distinguished Professorship in Comparative Oncology Genetics at NC State University. ALS is supported by NCI R50 CA211249.

