## Supplementary methods, discussion, tables, and figures for "Genomic analysis reveals shared genes and pathways in human and canine angiosarcoma"

*Sample collection*
Samples were collected from dogs referred for treatment at the University of Minnesota (UMN) Veterinary Medical Center, from samples submitted to the Modiano laboratory for diagnostic assessment and/or use in research (n = 34), from cases seen at the North Carolina State University (NCSU) Veterinary Hospital (n = 8), or from diagnostic samples sent to Antech Diagnostics (n = 5). Cases where date of diagnosis was known (n = 36), were diagnosed between 2000 and 2014. Tumor samples were either frozen (n = 17), or formalin-fixed and paraffin-embedded (FFPE, n = 30, **Table S1**). Procedures involving animal use were approved by the Institutional Animal Care and Use Committees at the Broad Institute, UMN, or NCSU.

*DNA extraction*
FFPE tumor samples were macrodissected by microtome to select for regions of high tumor cell density, and tumor DNA was prepared using the Qiagen QIAamp DNA FFPE Tissue Kit. DNA from frozen tumor tissue and germline DNA from whole blood were prepared using the Qiagen DNeasy Blood and Tissue Kit. The DNA was then eluted into nuclease-free purified water. DNA concentration was determined using the NanoDrop microvolume spectrophotometer (ThermoFisher Scientific) and/or the Quant-iT PicoGreen system (Invitrogen).

*Library construction*
1μg of DNA from each tumor and normal sample was diluted in Tris-EDTA (TE) buffer for sonic fragmentation. Samples were fragmented to a target size of 500bp using a Covaris ultrasonicator (Covaris). Fragments were cleaned and subject to size selection using Agencourt AMPure XP magnetic beads (Beckman Coulter). Select samples were visualized using the Agilent BioAnalyzer to check the distribution of fragment sizes. The Kapa Hyper Prep Kit was used for library construction (Kapa Biosystems). Briefly, size-selected DNA fragments were subject to end-repair and A-Tailing reactions, followed by attachment of adaptors in preparation for molecular barcoding. Fragments were purified using Agencourt AMPure XP magnetic beads between reactions. NEBNext Multiplex Oligos for Illumina (New England BioLabs) were then used to barcode the individual libraries following product guidelines.

*Amplification of samples*
The 66 overamplified samples had significantly fewer mutations than the 28 samples amplified using the recommended number of PCR cycles (mean mutational burden per tumor/normal pair = 21.4 vs 31.3, $p = 0.02$) and significantly lower estimated library size (number of unique molecules; $p < 0.001$; **Table S2**) consistent with a lower library complexity in the overamplified samples. Thus, in the overamplified samples, we are likely underpowered to discover variants, leading to a more conservative set of mutations. We note that all significantly mutated genes discovered during data analysis are mutated in both standard and overamplified samples, with the sole exception of *ATP5PD*, which was only found in the overamplified samples (**Figure S2**).

*Exome capture*

Custom blocking oligonucleotides were designed complementary to the barcode sequences and synthesized by Integrated DNA Technologies (IDT). Briefly, the amplified libraries were incubated with the SeqCap EZ probes, blocking oligos, and Roche Developer Reagent (in place of human Cot-1 DNA), for 60 hours in a thermocycler (Eppendorf). Captured DNA was then recovered and washed using SeqCap-EZ Capture Beads. The captured library was amplified via ligation-mediated (LM)-PCR for the recommended 14 cycles, and the amplified captured library washed using AMPure XP magnetic beads. qPCR using primers for specific targeted regions and a negative control region that was not targeted was performed to test enrichment of a subset of the captured libraries for using the LightCycler 480 instrument (Roche).

*Somatic variant calling*
We called variants in the GATK3 MuTect2 and GATK4 Mutect2 versions, with the addition of the *--dontUseSoftClippedBases* option. Prior to using this setting, we saw a large number of artifactual indels being called in our FFPE samples. These indels were being called with the only support for the variant being the ends of soft-clipped reads. A similar artifact has been reported in WGA TCGA data (1). We found no significant difference in the total number of mutations ($p_{t\text{-}test}$ = 0.76) or percent of indels (p = 0.95) between frozen (n = 17) and FFPE (n =30) samples in the final somatic mutation call set.

Variants for the panel of normals were called as recommended in the GATK3 workflow, using *--artifactDetectionMode* in MuTect2, and the calls from the normal samples were merged using CombineVariants, keeping any variant that was called in two or more dogs. All preprocessing was performed using GATK version 3.6.0 (2). BQSR was performed using a set of 19,112,082 known canine SNP positions drawn from multiple sources, including SNPs discovered by the Lindblad-Toh (3,4) and Axelsson labs (5), those included on the Affymetrix Axiom Canine HD array, and those contained in the DoGSD database (6).

Using GATK4, we again used the default Mutect2 parameters, with the addition of the *--dontUseSoftClippedBases* option. We then applied the FilterMutectCalls tool to this set of variant calls, using default cutoffs, with the exception of increasing the stringency of the median read position filter to ten, and specifying a unique alternate allele read count of four. The FilterByOrientationBias tool (labeled "experimental") was also applied, using the "G,T" setting (for oxidation artifacts) for all samples, and the "C,T" setting for FFPE-preserved samples.

*Somatic copy number aberration calling*
Somatic copy number alterations in tumors compared to the matched normal were called in the exome data using VarScan2 (7) followed by circular binary segmentation to translate intensity measurements into regions. Recurrent SCNAs were then identified using Gistic2 (8) using default options and a cutoff threshold of 10000 for "max seg".

*RNA-sequencing*
Total RNA was isolated from tissue samples using the TriPure Isolation Reagent (Roche Applied Science), and the RNeasy Mini Kit (Qiagen) was used for clean-up according to the manufacturer's instructions. Briefly, the TruSeq RNA sample preparation kit (Illumina) and a HiSeq 2000 or 2500

sequencing system (Illumina) were used to generate Illumina sequencing libraries. Each sample was sequenced to a targeted depth of approximately 20 – 80 million paired-end reads with mate-pair distance of 50 bp. Primary analysis and demultiplexing were performed using CASAVA software version 1.8.2 (Illumina) to verify the quality of the sequence data. The end result of the CASAVA workflow was demultiplexed into FASTQ files for analysis. Bioanalyzer quality control and RNA-seq were performed at the University of Minnesota Genomics Center (UMGC) or at the Broad Institute.

*Comparative pathway analysis*

For the comparative pathway analysis, gene names and Ensembl IDs were mapped to known gene IDs within the DAVID tool. The 1092 total canine mutations mapped to 951 human gene IDs in the DAVID tool. The 1958 total human genes mapped to 1934 gene IDs. The union of all mutated genes in both cohorts (n=2929) mapped to 2617 IDs.

*Construction of lollipop plots*

Canine variants were lifted over to the human genome (hg19) using the UCSC LiftOver tool (9). Plots were created using the MutationMapper function at CBioPortal. Six canine TP53 variants were not plotted due to mismatch of the reference allele in the canine and human genomes.

### Supplementary discussion

*Significantly mutated genes in the canine cohort*

The significantly mutated genes in our canine cohort which were less frequently mutated still may give us important insight into disease pathophysiology. *RASA1* is a negative regulator of the RAS and MAPK pathways, and plays an important role in vascular formation(10,11). Germline *RASA1* mutations can cause capillary malformation - arteriovenous malformation syndrome(12). Somatic mutations in this gene have been found in a subset of human basal cell carcinomas, and expression has been correlated with survival in invasive ductal breast carcinomas and hepatocellular carcinomas(13,14). Three out of four of the mutations in this gene in our dataset are nonsense mutations. In the Angiosarcoma Project data, one case has a missense mutation and one case a nonsense mutation in *RASA1*. *ORC1* is part of the DNA replication complex(15). *ARPC1A* plays an important role in regulating the actin cytoskeleton, which functions in the migration and invasiveness of pancreatic carcinoma cells(16). *ATP5H* is a mitochondrial ATP synthase(15). One study found that ATP5H interacts with PLCB1, which promotes cellular proliferation(17), while another study found that loss of ATP5H activity led to a stem cell-like phenotype, invasiveness, and resistance to therapy(18). It is notable that this gene was also significantly mutated in golden retriever B-cell lymphomas in one of our earlier exome sequencing studies(19).

*Mutational burden by tumor location*

In the human data, mutation rates are significantly different between different tumor locations, with head and neck angiosarcomas having a much higher mutational burden, as reported by Painter, et al (20). It is possible that the higher mutational burden in these tumors is due to UV exposure. It would be interesting to compare these findings to canine cutaneous and subcutaneous angiosarcomas, which were not included in this study. In the canine data, no significant difference was found between mutational burden by tumor location, however, all included cases were visceral tumors. It makes sense that the golden retrievers have a lower overall somatic mutation burden than humans, as they likely have a much higher germline risk burden, given the high incidence of angiosarcoma in the breed.

*RNA-seq survey of angiosarcoma in multiple breeds*

We took advantage of the broader cohort of canine angiosarcoma cases in the RNA-seq data to look at the distribution of SMGs across multiple breeds, and found that frequent *TP53* and *PIK3CA* mutations are not unique to the golden retriever genetic background, but are also found in other breeds, as reported by Wang, *et al (21)*.

*Comparative pathway analysis*

The most highly enriched category in both the dog and human filtered data was glycoproteins, likely because this broad category actually encompasses many of the protein families that are noted as enriched on their own. Glycoproteins are a large category of proteins which have been post-translationally modified to carry a carbohydrate group (22), and play a role in numerous cellular processes, including playing an important role in the extracellular signalling, through membrane receptors and secreted proteins.

While *PLCG1* has been reported as frequently mutated in human angiosarcomas, particularly in secondary tumors (23), none of the cases with *PLCG1* mutations in the current Angiosarcoma Project had previous radiation therapy (two had radiation as part of their current treatment for angiosarcoma), perhaps suggesting that mutations in this gene are not as rare in primary angiosarcomas as previously thought.

13. Liu Y, Liu T, Sun Q, Niu M, Jiang Y, Pang D. Downregulation of Ras GTPase‑activating protein 1 is associated with poor survival of breast invasive ductal carcinoma patients. Oncol Rep.

2015;33:119–24.

20. Painter C, Jain E, Dunphy M, Anastasio E, McGillicuddy M, Stoddard R, Thomas B, Balch Sara, Anderka K, Larkin K, Lennon N, Chen Y-L, Zimmer A, Baker EO, Maiwald S, Lapan J, Hornick JL, Raut C, Demetri G, Lander ES, Golub T. High mutation burden and response to immune checkpoint inhibitors in angiosarcomas of the scalp and face. 2018.

21. Wang G, Wu M, Maloneyhuss MA, Wojcik J, Durham AC, Mason NJ, et al. Actionable mutations in canine hemangiosarcoma. PLoS One. 2017;12:e0188667.

22. Wilkinson A, McNaught A. IUPAC Compendium of Chemical Terminology, (the" Gold Book"). International Union of Pure and Applied Chemistry. 1997;

23. Murali R, Chandramohan R, Möller I, Scholz SL, Berger M, Huberman K, et al. Targeted massively parallel sequencing of angiosarcomas reveals frequent activation of the mitogen activated protein kinase pathway. Oncotarget. 2015;6:36041–52.

**Supplementary Tables and Figures**
**Genomic analysis reveals shared genes and pathways in human and canine angiosarcoma**

**Table S1 - Cohort metadata**

| Case ID | Tissue preservation | Location | Nonsynonymous mutations | Sex | Age |
|---|---|---|---|---|---|
| 1 | fixed | liver | 15 | F | 10 |
| 2 | frozen | spleen | 22 | M | 9 |
| 3 | fixed | heart | 35 | F | 11 |
| 4 | frozen | spleen | 26 | F | 10 |
| 5 | fixed | liver | 30 | M | 11 |
| 6 | frozen | spleen | 22 | M | 10 |
| 7 | fixed | liver | 17 | M | 6 |
| 8 | fixed | heart | 31 | F | 5 |
| 9 | frozen | spleen | 17 | F | 11 |
| 10 | fixed | heart | 10 | F | 12 |
| 11 | frozen | spleen | 26 | M | 7 |
| 12 | fixed | heart | 22 | M | 10 |
| 13 | fixed | spleen | 13 | F | 8 |
| 14 | fixed | spleen | 11 | F | 11 |
| 15 | fixed | heart | 24 | F | 15 |
| 16 | frozen | spleen | 37 | M | 10 |
| 17 | frozen | heart | 33 | F | 8 |
| 18 | frozen | spleen | 13 | M | 9 |
| 19 | frozen | spleen | 17 | M | 3 |
| 20 | frozen | heart | 21 | M | 9 |
| 21 | fixed | heart | 64 | F | 8 |
| 22 | fixed | heart | 35 | M | 12 |
| 23 | frozen | spleen | 5 | F | 11 |
| 24 | fixed | liver | 37 | F | 10 |
| 25 | fixed | liver | 27 | M | 12 |
| 26 | frozen | spleen | 3 | F | 14 |
| 27 | frozen | spleen | 41 | F | 9 |
| 28 | frozen | heart | 45 | M | 9 |
| 29 | frozen | spleen | 26 | M | 13 |
| 30 | fixed | spleen | 3 | M | NA |
| 31 | fixed | spleen | 43 | M | NA |
| 32 | fixed | spleen | 22 | F | NA |

| | | | | | |
|---|---|---|---|---|---|
| 33 | fixed | spleen | 7 | M | NA |
| 34 | fixed | spleen | 13 | M | NA |
| 35 | fixed | heart | 46 | F | 11 |
| 36 | fixed | heart | 16 | F | 9 |
| 37 | fixed | heart | 149 | F | 15 |
| 38 | fixed | spleen | 13 | M | NA |
| 39 | fixed | heart | 33 | F | NA |
| 40 | fixed | heart | 16 | F | NA |
| 41 | frozen | heart | 30 | M | NA |
| 42 | fixed | liver | 23 | M | NA |
| 43 | fixed | spleen | 46 | F | NA |
| 44 | frozen | spleen | 14 | F | 4 |
| 45 | fixed | liver | 22 | F | 5 |
| 46 | fixed | liver | 41 | F | 13 |
| 47 | fixed | unknown, likely heart | 1 | F | 11 |

**Table S2 - Library preparation, amplification, and complexity**

| Case ID | Tumor/ Normal | Sequencing depth | Input DNA | Overamplified | Tissue preservation | Nonsynonymous mutations | Library size |
|---|---|---|---|---|---|---|---|
| 1 | Normal | 49 | 1000 | Yes | EDTA | | 134561459 |
| 1 | Tumor | 53 | 1000 | Yes | fixed | 15 | 38155922 |
| 2 | Normal | 71 | 1000 | Yes | EDTA | | 226048914 |
| 2 | Tumor | 91 | 1000 | Yes | frozen | 22 | 136254048 |
| 3 | Normal | 73 | 1000 | Yes | EDTA | | 134978503 |
| 3 | Tumor | 26 | 1000 | Yes | fixed | 35 | 20412528 |
| 4 | Normal | 61 | 1000 | Yes | EDTA | | 231846809 |
| 4 | Tumor | 99 | 1000 | Yes | frozen | 26 | 103917348 |
| 5 | Normal | 51 | 1000 | Yes | EDTA | | 370312244 |
| 5 | Tumor | 41 | 1000 | Yes | fixed | 30 | 32017163 |
| 6 | Normal | 63 | 1000 | Yes | EDTA | | 225490375 |
| 6 | Tumor | 84 | 1000 | Yes | frozen | 22 | 130755435 |
| 7 | Normal | 53 | 1000 | Yes | EDTA | | 269484482 |
| 7 | Tumor | 89 | 1000 | Yes | fixed | 17 | 73256644 |
| 8 | Normal | 80 | 1000 | Yes | EDTA | | 268801501 |
| 8 | Tumor | 75 | 1000 | Yes | fixed | 31 | 81740488 |

| | | | | | | | |
|---|---|---|---|---|---|---|---|
| 9 | Normal | 62 | 1000 | Yes | EDTA | | 281443846 |
| 9 | Tumor | 101 | 1000 | Yes | frozen | 17 | 210031996 |
| 10 | Normal | 83 | 1000 | No | EDTA | | 355514908 |
| 10 | Tumor | 78 | 1000 | No | fixed | 10 | 69807014 |
| 11 | Normal | 68 | NA | No | EDTA | | 277032267 |
| 11 | Tumor | 126 | 1000 | No | frozen | 26 | 324015250 |
| 12 | Normal | 62 | 1000 | Yes | EDTA | | 221713689 |
| 12 | Tumor | 55 | 1000 | Yes | fixed | 22 | 49450048 |
| 13 | Normal | 43 | 1000 | Yes | EDTA | | 203071580 |
| 13 | Tumor | 91 | 1000 | Yes | fixed | 13 | 144056227 |
| 14 | Normal | 58 | 1000 | Yes | EDTA | | 177387584 |
| 14 | Tumor | 61 | 1000 | Yes | fixed | 11 | 65437960 |
| 15 | Normal | 75 | 1000 | Yes | EDTA | | 127216800 |
| 15 | Tumor | 56 | 1000 | Yes | fixed | 24 | 40991888 |
| 16 | Normal | 73 | 1000 | Yes | EDTA | | 141452611 |
| 16 | Tumor | 56 | 1000 | Yes | frozen | 37 | 41118918 |
| 17 | Normal | 62 | 1000 | No | EDTA | | 223257784 |
| 17 | Tumor | 114 | NA | No | frozen | 33 | 147787071 |
| 18 | Normal | 53 | 1000 | Yes | EDTA | | 171084585 |
| 18 | Tumor | 78 | 1000 | Yes | frozen | 13 | 97513922 |
| 19 | Normal | 69 | 1000 | Yes | EDTA | | 129594138 |
| 19 | Tumor | 75 | 1000 | Yes | frozen | 17 | 91271441 |
| 19 | Normal | 67 | 1000 | Yes | EDTA | | 110209804 |
| 20 | Tumor | 91 | 1000 | Yes | frozen | 21 | 161197314 |
| 21 | Normal | 64 | 1000 | No | EDTA | | 293332856 |
| 21 | Tumor | 144 | 309* | No | fixed | 64 | 161501907 |
| 22 | Normal | 62 | 1000 | Yes | EDTA | | 123858089 |
| 22 | Tumor | 57 | 1000 | Yes | fixed | 35 | 42916453 |
| 23 | Normal | 54 | 1000 | Yes | EDTA | | 214634405 |
| 23 | Tumor | 84 | 1000 | Yes | frozen | 5 | 163422686 |
| 24 | Normal | 51 | 1000 | Yes | EDTA | | 130312170 |
| 24 | Tumor | 56 | 1000 | Yes | fixed | 37 | 40995053 |
| 25 | Normal | 62 | 1000 | No | EDTA | | 232499575 |
| 25 | Tumor | 46 | 1000 | Yes | fixed | 27 | 34254483 |
| 26 | Normal | 74 | 1000 | No | EDTA | | 159151355 |
| 26 | Tumor | 93 | 381* | No | frozen | 3 | 128242059 |
| 27 | Normal | 74 | 1000 | Yes | EDTA | | 100619527 |

| | | | | | | | |
|---|---|---|---|---|---|---|---|
| 27 | Tumor | 96 | 1000 | Yes | frozen | 41 | 135472727 |
| 28 | Normal | 90 | 1000 | No | EDTA | | 229798708 |
| 28 | Tumor | 109 | 469* | No | frozen | 45 | 196472240 |
| 29 | Normal | 52 | 1000 | No | EDTA | | 280852929 |
| 29 | Tumor | 125 | 425* | No | frozen | 26 | 316081507 |
| 30 | Normal | 69 | 1000 | Yes | EDTA | | 131692351 |
| 30 | Tumor | 85 | 1000 | Yes | fixed | 3 | 100942519 |
| 31 | Normal | 50 | 1000 | Yes | EDTA | | 118651337 |
| 31 | Tumor | 75 | 1000 | Yes | fixed | 43 | 48558498 |
| 32 | Normal | 59 | 1000 | Yes | EDTA | | 137223693 |
| 32 | Tumor | 40 | 1000 | Yes | fixed | 22 | 25433630 |
| 33 | Normal | 65 | 1000 | Yes | EDTA | | 116541031 |
| 33 | Tumor | 54 | 1000 | Yes | fixed | 7 | 63455690 |
| 34 | Normal | 60 | 1000 | Yes | EDTA | | 89817007 |
| 34 | Tumor | 74 | 1000 | Yes | fixed | 13 | 44486076 |
| 35 | Normal | 51 | 1000 | No | EDTA | | 210786142 |
| 35 | Tumor | 114 | 295* | No | fixed | 46 | 163041517 |
| 36 | Normal | 27 | 1000 | Yes | EDTA | | 58556571 |
| 36 | Tumor | 45 | 1000 | Yes | fixed | 16 | 56260934 |
| 37 | Normal | 54 | 1000 | No | EDTA | | 98731048 |
| 37 | Tumor | 128 | 871* | No | fixed | 149 | 127593218 |
| 37 | Normal | 67 | 1000 | Yes | EDTA | | 87139774 |
| 38 | Tumor | 45 | 1000 | Yes | fixed | 13 | 31958634 |
| 39 | Normal | 95 | NA | No | EDTA | | 171064700 |
| 39 | Tumor | 93 | 355* | No | fixed | 33 | 88831176 |
| 40 | Normal | 54 | 1000 | Yes | EDTA | | 65023743 |
| 40 | Tumor | 18 | 1000 | Yes | fixed | 16 | 16470714 |
| 41 | Normal | 59 | 1000 | No | EDTA | | 173375722 |
| 41 | Tumor | 97 | 480* | No | frozen | 30 | 119131774 |
| 42 | Normal | 98 | 1000 | No | EDTA | | 190927431 |
| 42 | Tumor | 108 | 941* | No | fixed | 23 | 120828418 |
| 43 | Normal | 51 | 1000 | Yes | EDTA | | 90620114 |
| 43 | Tumor | 158 | 373* | No | fixed | 46 | 206945827 |
| 44 | Normal | 60 | 1000 | Yes | EDTA | | 50917493 |
| 44 | Tumor | 42 | 1000 | Yes | frozen | 14 | 32736117 |
| 45 | Normal | 88 | 1000 | No | EDTA | | 288287484 |
| 45 | Tumor | 108 | 292* | No | fixed | 22 | 108587264 |

| | | | | | | | |
|---|---|---|---|---|---|---|---|
| 46 | Normal | 33 | 1000 | Yes | EDTA | | 27281614 |
| 46 | Tumor | 40 | 1000 | Yes | fixed | 41 | 27533702 |
| 47 | Normal | 57 | 1000 | Yes | EDTA | | 48624834 |
| 47 | Tumor | 16 | 1000 | Yes | fixed | 1 | 14145086 |

DNA input amount, whether library was overamplified, and estimated library size (unique molecules) per library.
* estimated input amount.

**Table S3 - COSMIC Cancer Gene Census genes**

| **Gene** | **# samples** |
|---|---|
| ***TP53*** | 28 |
| ***PIK3CA*** | 14 |
| ***PIK3R1*** | 4 |
| *CHD4* | 3 |
| *CACNA1D* | 2 |
| *NRAS* | 2 |
| *PTEN* | 2 |
| *AR* | 1 |
| *ARID1A* | 1 |
| *ATR* | 1 |
| *ATRX* | 1 |
| *BRAF* | 1 |
| *BRCA2* | 1 |
| *CASP8* | 1 |
| *DICER1* | 1 |
| *ERBB4* | 1 |
| *FGFR3* | 1 |
| *FLT3* | 1 |
| *IL6ST* | 1 |
| *JAK1* | 1 |
| *KDR* | 1 |
| *KEAP1* | 1 |
| *KMT2D* | 1 |
| *LRP1B* | 1 |
| *MAP3K13* | 1 |
| *MTOR* | 1 |
| *NCOR1* | 1 |

| | |
|---|---|
| *NFE2L2* | 1 |
| *NTRK3* | 1 |
| *PLCG1* | 1 |
| *RNF43* | 1 |
| *TERT* | 1 |
| *ZFHX3* | 1 |

Genes mutated at least once in canine exome data, annotated as likely cancer drivers in the COSMIC Cancer Gene Census. Significantly mutated genes bolded.

**Table S4 - RNA-seq validation**

| ID | Gene | Exome variant | Tissue preservation | Mutation type | VAF | Validated | RNA-seq pileup |
|---|---|---|---|---|---|---|---|
| 17 | ***ARPC1A*** | chr6:10238294:G:A | frozen | Nonsense | 0.17 | No | 109\|0:0:0:0:0:0:0:0:0 |
| 36 | ***ARPC1A*** | chr6:10232318:A:C | fixed | Missense | 0.16 | Yes | 283\|0:0:36:0:0:1:0:0:0 |
| 21 | ***ORC1*** | chr15:8982528:A:C | fixed | Missense | 0.05 | No | 9\|0:0:0:0:0:0:0:0:0 |
| 4 | ***PIK3CA*** | chr34:12675674:A:T | frozen | Missense | 0.25 | No | 67\|0:0:0:11:0:0:0:0:0 |
| 18 | ***PIK3CA*** | chr34:12651933:G:A | frozen | Missense | 0.06 | No | 41\|0:1:0:0:0:0:0:0:0 |
| 27 | ***PIK3CA*** | chr34:12675674:A:G | fixed | Missense | 0.23 | No | 38\|0:0:0:1:0:1:0:0:0 |
| 28 | ***PIK3CA*** | chr34:12675674:A:T | fixed | Missense | 0.19 | Yes | 24\|0:3:0:0:0:0:0:0:0 |
| 29 | ***PIK3CA*** | chr34:12675674:A:G | fixed | Missense | 0.11 | Yes | 29\|0:0:0:11:0:0:0:0:0 |
| 11 | ***PIK3CA*** | chr34:12675666:T:G | frozen | Missense | 0.11 | Yes | 27\|0:0:0:13:0:0:0:0:0 |
| 36 | ***PIK3CA*** | chr34:12651893:G:A | fixed | Missense | 0.15 | Yes | 35\|10:0:0:0:0:0:0:0:0 |
| 21 | ***PIK3R1*** | chr2:53522035:A:G | fixed | Missense | 0.17 | Yes | 41\|0:0:0:5:0:0:0:0:0 |
| 18 | ***RASA1*** | chr3:21093342:G:A | frozen | Nonsense | 0.11 | No | 43\|0:1:0:0:0:0:0:0:0 |
| 28 | ***RASA1*** | chr3:21108103:A:T | fixed | Nonsense | 0.13 | Yes | 29\|0:3:0:0:0:0:0:0:0 |
| 4 | ***TP53*** | chr5:32563923:C:T | frozen | Missense | 0.32 | No | 49\|0:0:0:0:0:0:0:0:0 |
| 17 | ***TP53*** | chr5:32563712:C:T | frozen | Missense | 0.47 | No | 59\|0:0:0:0:0:1:0:0:0 |
| 19 | ***TP53*** | chr5:32563056:G:A | frozen | Missense | 0.18 | Yes | 66\|14:0:0:0:0:0:0:0:0 |
| 20 | ***TP53*** | chr5:32563698:A:T | frozen | Missense | 0.26 | No | 40\|0:0:0:0:0:0:0:0:0 |
| 21 | ***TP53*** | chr5:32563404:C:T | fixed | Missense | 0.27 | Yes | 125\|0:40:0:1:0:0:0:0:0 |
| 21 | ***TP53*** | chr5:32564616:A:G | fixed | Missense | 0.19 | Yes | 95\|0:0:0:35:0:0:0:0:0 |
| 22 | ***TP53*** | chr5:32563049:C:A | fixed | Missense | 0.1 | No | 194\|2:0:0:0:0:0:0:0:0 |
| 22 | ***TP53*** | chr5:32563400:C:T | fixed | Missense | 0.32 | No | 146\|0:0:0:0:0:0:0:0:0 |
| 24 | ***TP53*** | chr5:32563401:C:T | fixed | Missense | 0.27 | Yes | 129\|0:33:0:0:0:0:0:0:0 |
| 24 | ***TP53*** | chr5:32563916:C:T | fixed | Missense | 0.21 | Yes | 105\|0:27:0:1:0:0:0:0:0 |
| 29 | ***TP53*** | chr5:32563959:CCATAG:C | fixed | Frame shift | 0.09 | No | 25\|20:10:10:10:0:1:0:10:0 |
| 1 | ***TP53*** | chr5:32561417:CG:C | fixed | Frame shift | 0.17 | No | 82\|0:0:0:15:0:2:0:14:0 |

| | | | | | | | |
|---|---|---|---|---|---|---|---|
| **1** | ***TP53*** | chr5:32563056:G:A | fixed | Missense | 0.07 | Yes | 52\|14:0:0:0:0:2:0:0:0 |
| **11** | ***TP53*** | chr5:32564027:G:A | frozen | Missense | 0.42 | Yes | 147\|116:0:0:0:0:2:0:0:0 |
| **13** | ***TP53*** | chr5:32563400:C:T | frozen | Missense | 0.42 | Yes | 198\|0:148:0:0:0:0:0:0:0 |

**Table S5 - RNA-seq survey**

| Gene | Golden | Boxer | GSD | Lab | PWD | Keeshond | Other | Total |
|---|---|---|---|---|---|---|---|---|
| ***TP53*** | 18 | 0 | 1 | 0 | 2 | 1 | 1 | 23 |
| ***PIK3CA*** | 7 | 1 | 2 | 0 | 2 | 2 | 2 | 17 |
| ***PIK3R1*** | 2 | 0 | 0 | 1 | 1 | 0 | 1 | 5 |
| ***ARPC1A*** | 2 | 0 | 0 | 0 | 0 | 0 | 0 | 2 |

Survey of specific mutations identified in the exome sequencing in SMGs in RNA-seq data from canine angiosarcoma tumors in 43 golden retrievers and 31 dogs from other breeds.

**Table S6 DAVID functional clustering analysis of human genes**
See attached file.

**Table S7 DAVID functional clustering analysis of canine genes**
See attached file.

**Table S8 DAVID functional clustering analysis of genes overlapping between human and canine**
See attached file.

**Table S9 DAVID functional clustering analysis of the union of genes between human and canine**
See attached file.

**Table S10 - somatic copy number aberrations by oaCGH**

| Gene | % with gain | % with loss |
|---|---|---|
| ***TP53*** | 6 | 7 |
| ***PIK3CA*** | 10 | 4 |
| ***PIK3R1*** | 2 | 3 |
| ***ORC1*** | 7 | 2 |
| ***RASA1*** | 4 | 2 |
| ***ARPC1A*** | 2 | 0 |
| ***ATP5H*** | 0 | 0 |
| ***PTEN*** | 4 | 4 |

| | | |
|---|---|---|
| ***VEGFA*** | 19 | 3 |
| ***MYC*** | 9 | 0 |
| ***CDKN2A/B*** | 0 | 22 |
| ***KDR*** | 22 | 0 |
| ***KIT*** | 17 | 0 |

Somatic copy number aberrations in SMGs, as well as in select genes reported to be recurrently altered in canine and human literature, in a cohort of 66 angiosarcomas analyzed by oaCGH. (Thomas, *et al.*, manuscript in preparation)(23).

**Table S11 - comparison of top SCNAs in human Angiosarcoma Project data with canine oaCGH data**

| **Gene** | **CNA** | **# patients** | **% altered (human)** | **% gain (golden retriever)** | **% loss (golden retriever)** |
|---|---|---|---|---|---|
| ***OBSCN*** | DEL | 15 | 42 | 6 | 15 |
| ***KDR*** | **AMP** | **13** | **36** | **22** | **0** |
| ***DAXX*** | AMP | 12 | 33 | 6 | 0 |
| ***CD70*** | DEL | 12 | 33 | 13 | 2 |
| ***HOXA9*** | AMP | 11 | 31 | 10 | 6 |
| ***PPM1D*** | AMP | 10 | 28 | 3 | 2 |
| ***ZNF331*** | DEL | 10 | 28 | 2 | 0 |
| ***HOXA11*** | AMP | 10 | 28 | 9 | 6 |
| ***WWTR1*** | AMP | 10 | 28 | 2 | 3 |
| ***CD36*** | AMP | 10 | 28 | 6 | 0 |
| ***DDX5*** | AMP | 10 | 28 | 9 | 0 |
| ***TNFSF9*** | DEL | 8 | 22 | 13 | 2 |
| ***PTK2*** | AMP | 8 | 22 | 6 | 2 |
| ***MAP4K4*** | AMP | 8 | 22 | 2 | 9 |
| ***EGFL7*** | AMP | 7 | 19 | 7 | 19 |
| ***NDRG1*** | AMP | 7 | 19 | 10 | 3 |
| ***APH1A*** | AMP | 7 | 19 | 4 | 0 |
| ***KIT*** | **AMP** | **7** | **19** | **17** | **0** |

Percentage copy number gain or loss in oaCGH data from 69 golden retriever angiosarcomas compared with the most frequent SCNAs (affecting >15% of patients) occurring in known cancer genes in 48 tumor samples from The Angiosarcoma Project. SCNAs affecting >15% of patients in the same direction in both species bolded. (Thomas, *et al.*, manuscript in preparation)(23).

**Figure S1 - Canine exome sequencing workflow**

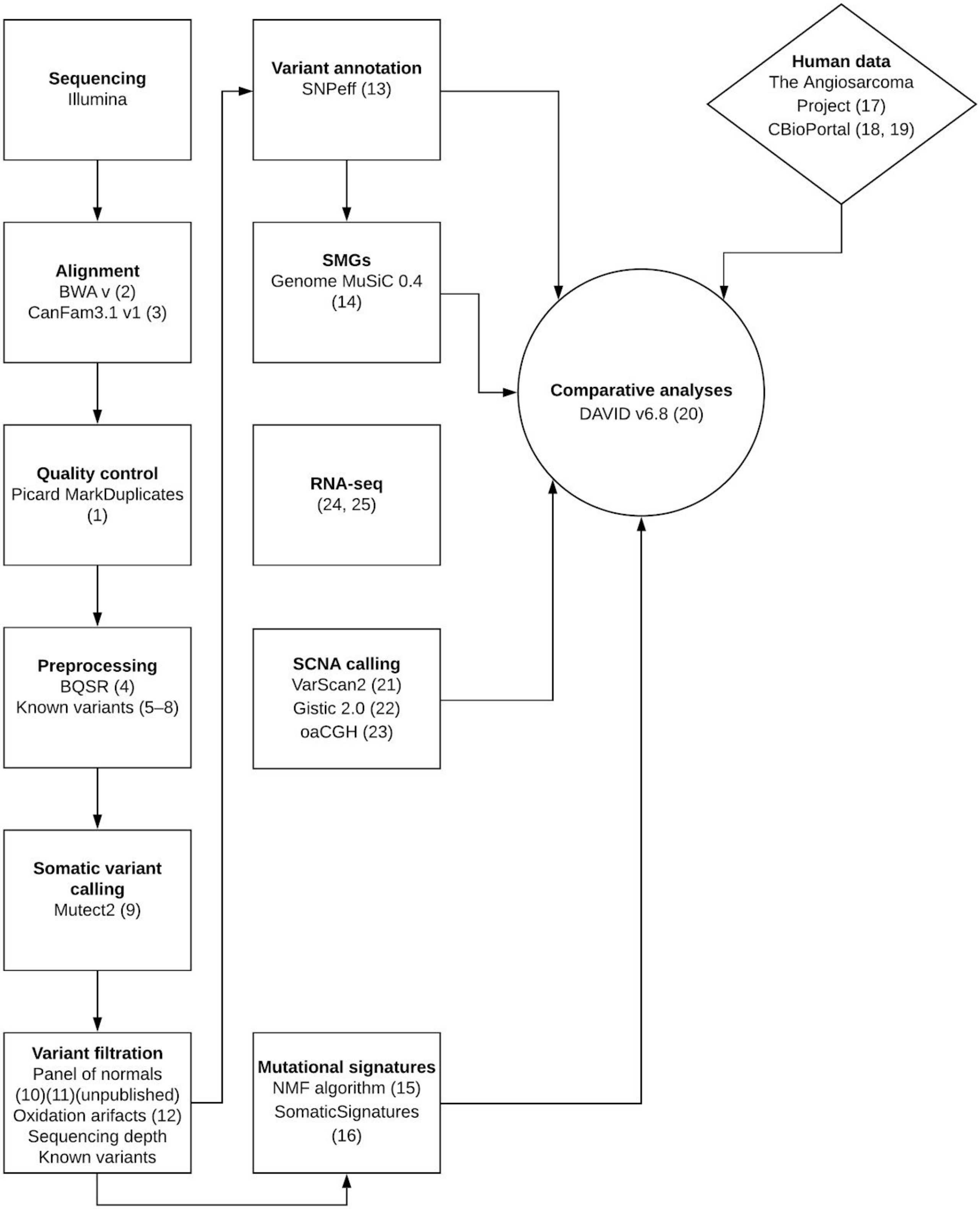

### Figure S2 - SMGs called in overamplified vs. normal libraries

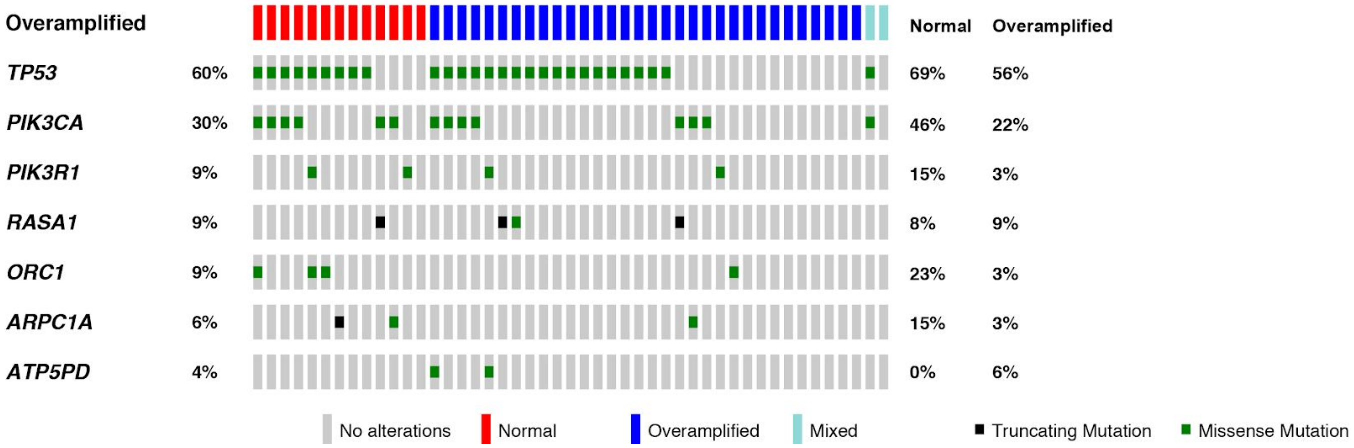

**Figure S3 - Mutational landscape of entire cohort**

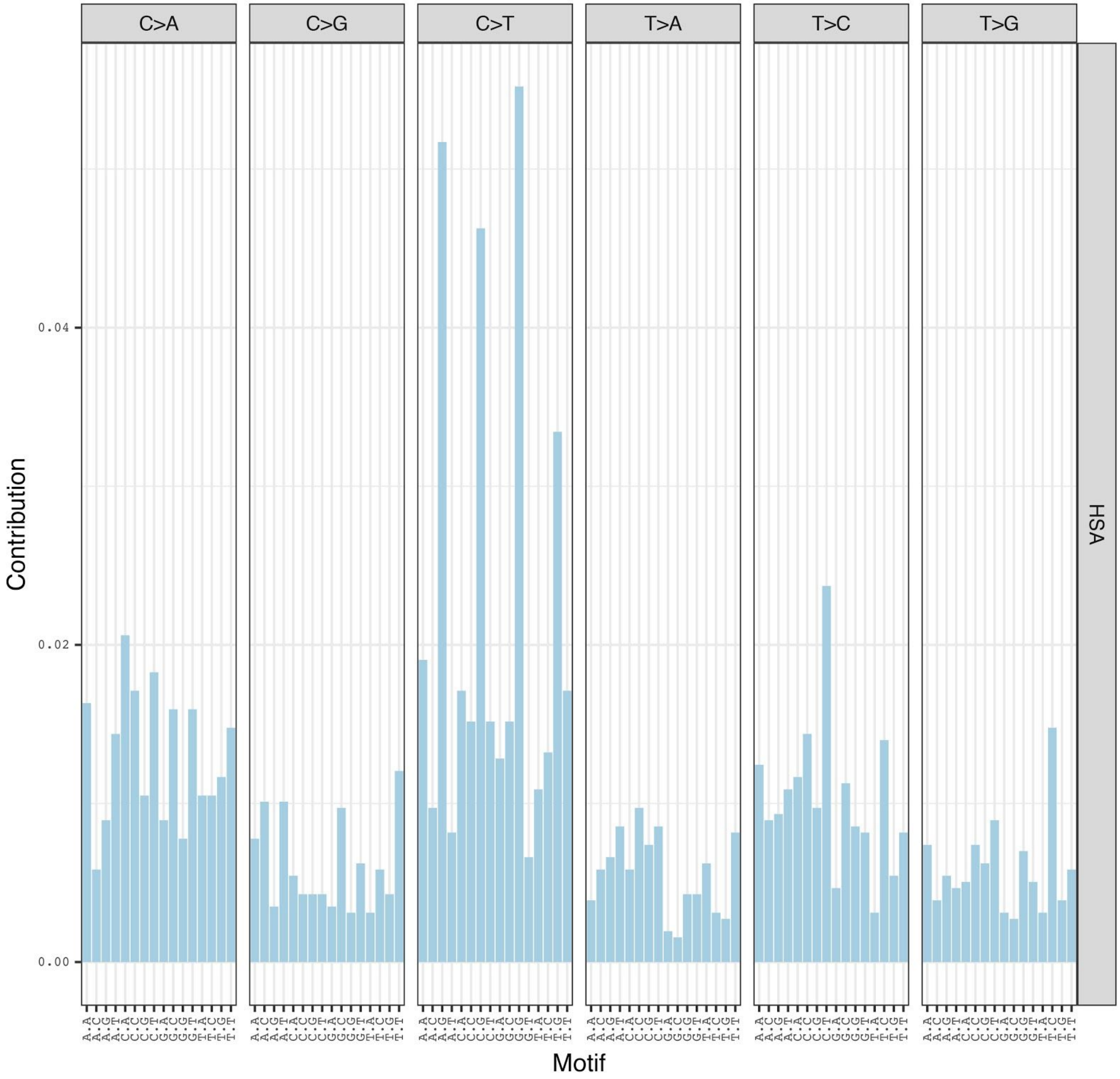


Mutational landscape of all canine angiosarcoma cases.

**Figure S4 - Mutational landscape: FFPE vs Frozen**

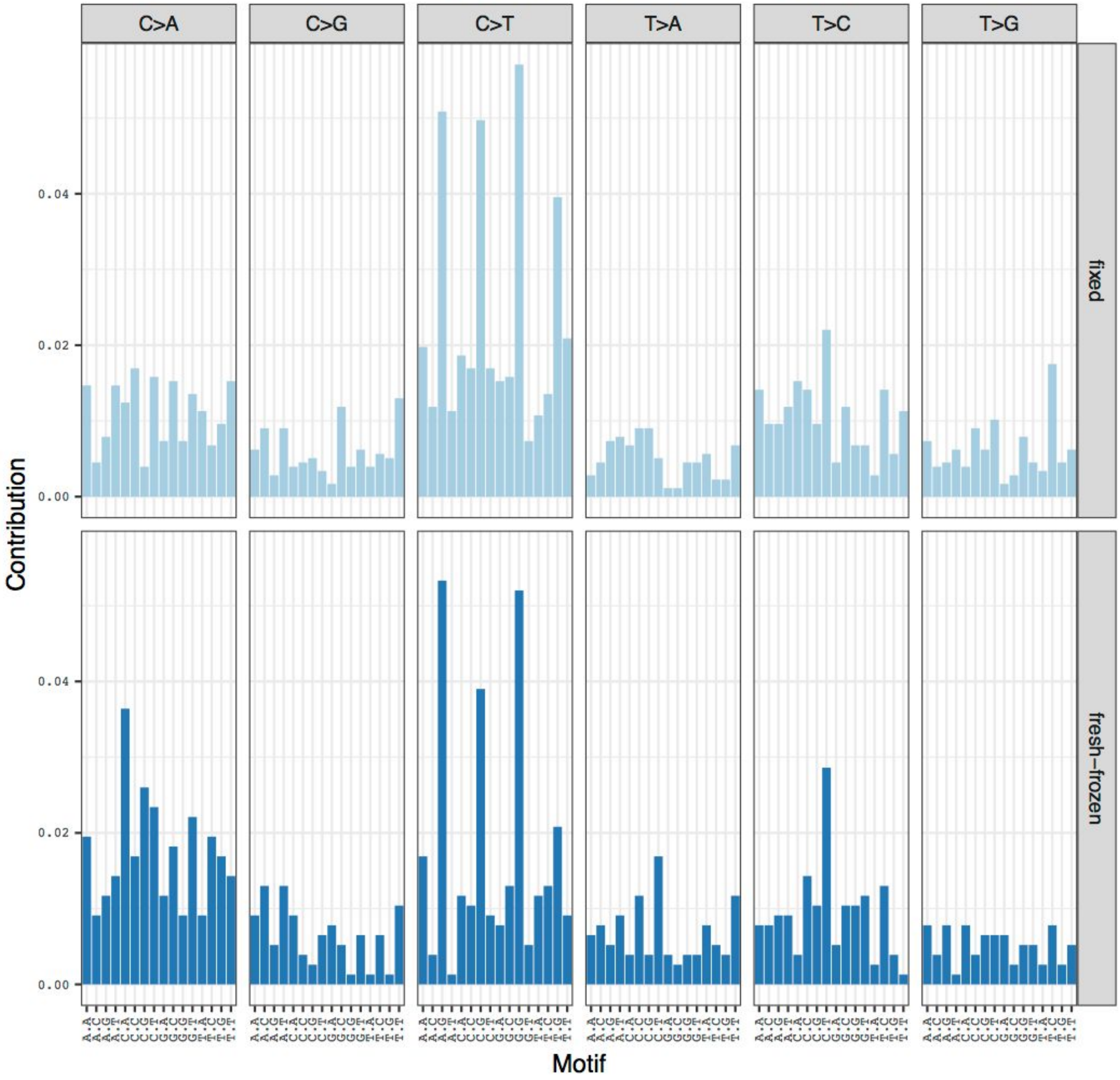


Mutational landscape of all canine cases, divided by tissue handling (fixed vs. frozen).

**Figure S5 - Mutational landscape: Tumor Location**

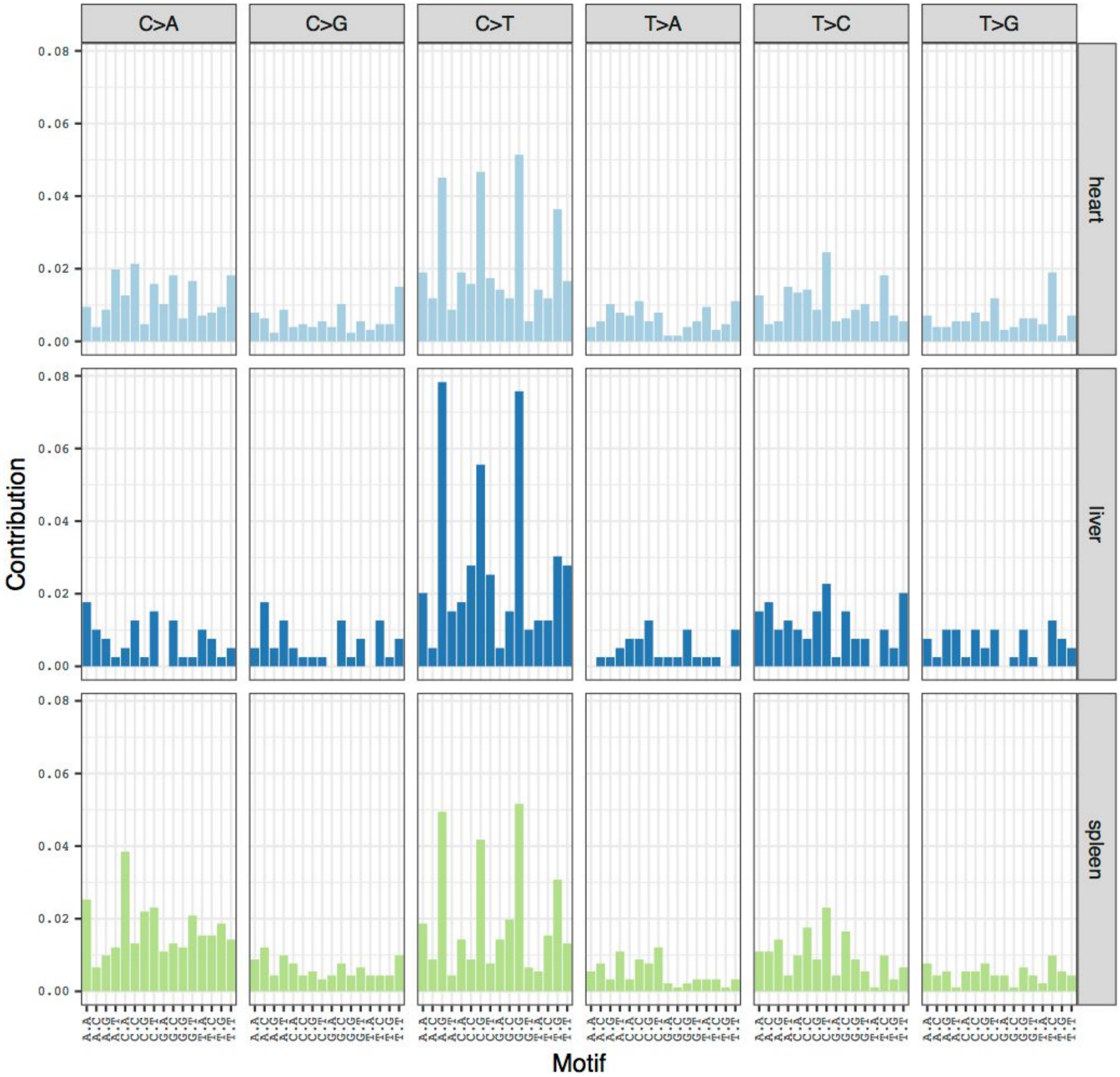


Mutational landscape of all canine cases, divided by tumor location (heart, liver, spleen).

**Figure S6 - FFPE vs Frozen exome SCNA data clustering**

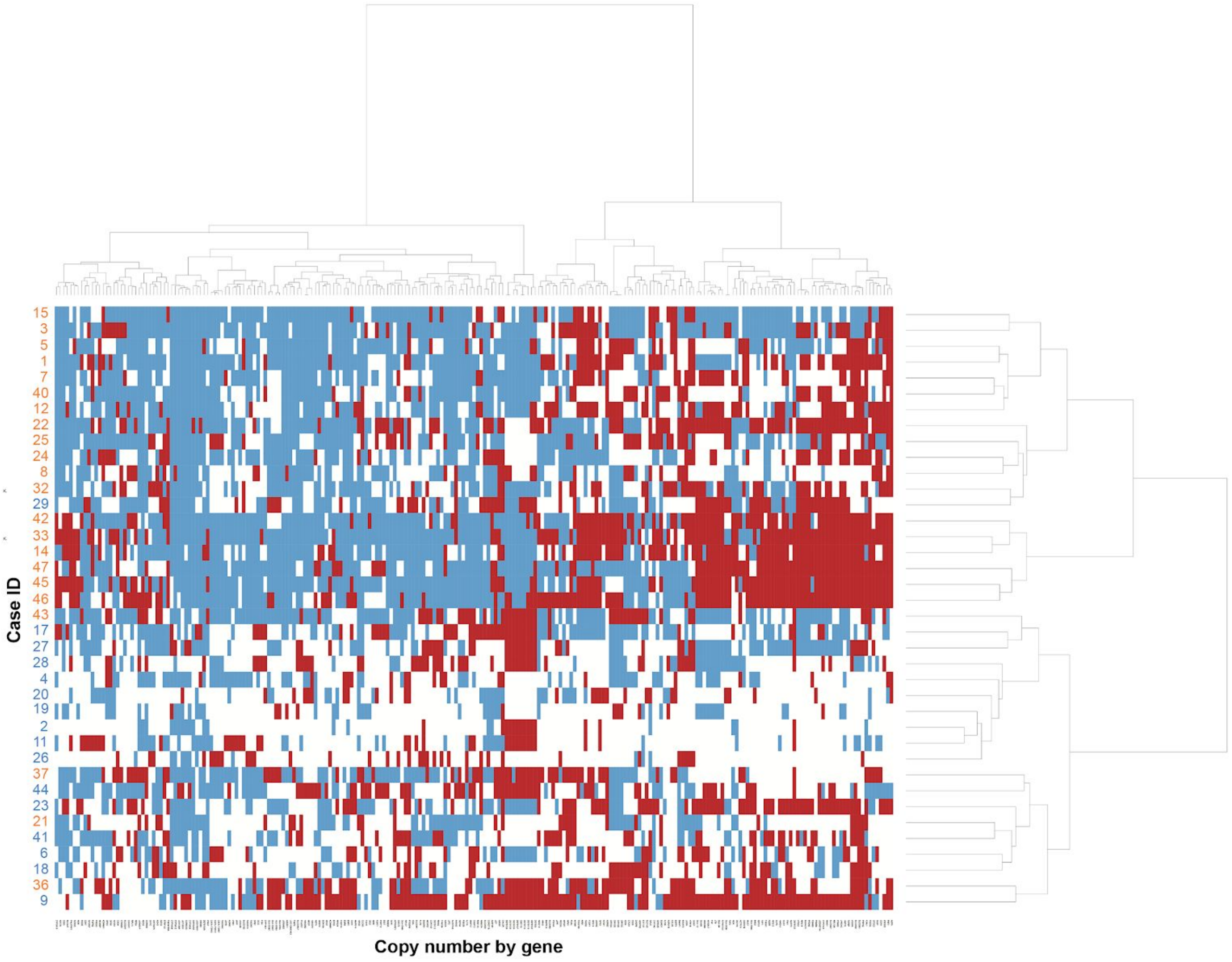


Unsupervised clustering of the VarScan2 copy number aberrations in the 38 canine angiosarcoma cases with fewer than 10,000 segments, showing clustering of amplifications (red) and deletions (red) per gene by sample. Case IDs show fixed (orange) or frozen (blue) tissue.

16. Gehring JS, Fischer B, Lawrence M, Huber W. SomaticSignatures: inferring mutational signatures from

single-nucleotide variants. Bioinformatics. 2015;31:3673–5.
